# HDAC1/2-mediated repression of Wnt receptor expression orients asymmetric division polarity in *C. elegans*

**DOI:** 10.64898/2026.02.23.707389

**Authors:** Mar Ferrando-Marco, Beatriz Garcia del Valle, Mark Hintze, Lucy Narunsky, Shuxiao Lin, Junyue Huang, Shannon Edwards, Michalis Barkoulas

## Abstract

Asymmetric cell division generates distinct daughter cells essential for tissue development, yet the mechanisms orienting division polarity within tissues remain incompletely understood. Here, we uncover a role for chromatin-mediated transcriptional repression in controlling polarity orientation during asymmetric division of *C. elegans* epidermal stem cells, known as the seam cells. We show that tissue-specific loss of the class I histone deacetylase *hda-1*, homologous to mammalian HDAC1/2, causes reversals in division polarity and reduces molecular asymmetry between daughter cells. Using Targeted DamID to profile HDA-1 genomic occupancy, we identify the Wnt receptors *lin-17*/Frizzled and *cam-1*/Ror as key targets. Single-molecule FISH reveals that these receptors display striking expression gradients along the body axis, with *cam-1* enriched anteriorly and *lin-17* posteriorly. In *hda-1* mutants, both receptors are upregulated so that differences between seam cells are flattened. Overexpression of either receptor alone is sufficient to reproduce the polarity reversals observed in *hda-1* mutants, and their co-overexpression produces additive defects. The polarity phenotype is independent of the canonical NuRD and SIN3 complexes, suggesting that HDA-1 acts through an alternative mechanism to regulate Wnt receptor expression. Our findings establish a direct link between histone deacetylase activity and Wnt receptor expression and support a model in which graded expression of Wnt receptors provides positional cues that orient asymmetric division polarity in response to Wnt signals.

## Introduction

The development of multicellular organisms relies on the generation of diverse cell types organised into functional tissues. Stem cells play a critical role in this process by undergoing asymmetric divisions, which generate daughter cells capable of differentiation as well as symmetric divisions that expand the stem cell population (Slack, 2018). In addition to generating daughter cells with distinct fates, asymmetric divisions must be precisely oriented within tissues. Proper orientation of polarity ensures that daughter cells are positioned appropriately and exposed to the correct extrinsic cues, as the directional specification of daughter cell fates determines their spatial organisation within tissues and establishes tissue architecture (Gómez-López et al., 2014; Morrison & Kimble, 2006; Williams & Fuchs, 2013). Defects in division orientation and polarity can disrupt tissue homeostasis and contribute to disease, including cancer, where loss of polarity and dysregulation of asymmetric division are frequently observed (Bajaj et al., 2020; Yamashita et al., 2010). Understanding the genes and pathways that regulate these fundamental processes is therefore crucial for elucidating mechanisms underlying both normal development and disease states.

*Caenorhabditis elegans* epidermal seam cells provide an excellent system for investigating stem cell behaviour and asymmetric division (Joshi et al., 2010). The seam cells are arranged in a linear pattern along the body axis (H0-H2, V1-V6, and T) and undergo both symmetric and asymmetric divisions throughout postembryonic development (Figure 1A). During asymmetric division, posterior daughters typically retain seam cell fate and express genes such as the GATA factors ELT-1, EGL-18, and ELT-6 that promote seam cell fate maintenance (Gorrepati et al., 2013; Koh & Rothman, 2001; Smith et al., 2005). Seam cells contribute most of the epidermal nuclei during post-embryonic development and hence produce the protective cuticle (Chisholm & Hsiao, 2012). Additionally, specific seam cell lineages (H2, V5, and T) generate neuronal precursors (Sulston & Horvitz, 1977). During the early L2 stage, a symmetric division expands the seam cell population from 10 to 16 cells, with both daughters retaining stem-like properties. At the end of larval development, these 16 cells undergo terminal differentiation and fuse to form a syncytium that produces the alae, specialised ridges in the cuticle (Altun & Hall, 2002). The highly reproducible nature of seam cell division patterns and their fixed anterior-posterior positioning make them an excellent system for detailed analysis of developmental defects at single-cell resolution (Katsanos et al., 2017).

**Figure 1.**
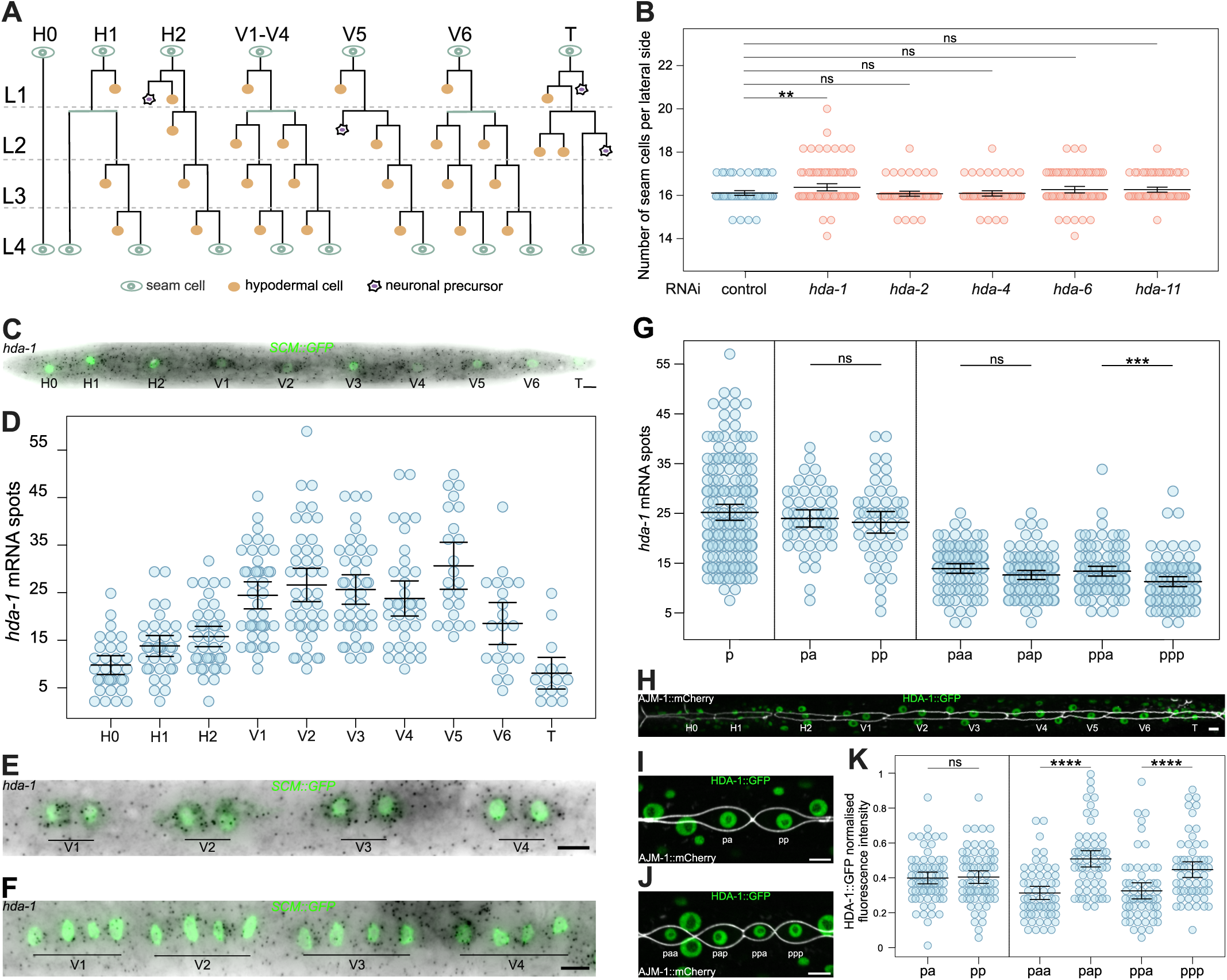
Analysis of *hda-1* expression at the mRNA and protein level in seam cells. **(A)** Schematic showing seam cell divisions from L1 to L4. Seam cells are labelled in green, hypodermal cells are labelled in orange and neuroblasts are labelled in purple. **(B)** Seam cell number counts upon RNAi targeting members of the HDAC family in *C. elegans*. Only downregulation of *hda-1* leads to a significant increase in the terminal seam cell number (p<0.01 with a two-tailed t-test, 65 ≤ n ≤ 96 animals per condition). **(C)** Representative *hda-1* smFISH image at late L1 stage. **(D)** Quantification of *hda-1* mRNA spots in all seam cell lineages at late L1 (21 ≤ n ≤ 45 cells per condition). **(E-F)** Representative *hda-1* smFISH images following the L2 symmetric division (E), and the L2 asymmetric division (F). **(G)** Quantification of *hda- 1* mRNA spots in V1-V4 seam cells before (p) and after the L2 symmetric division (pa, pp), or the L2 asymmetric division (paa, pap, ppa, ppp); n = 166 (p), n = 51 (pa, pp), n = 90 (paa, pap, ppa, ppp) animals per condition. **(H)** Representative image of HDA-1::GFP at late L1. **(I-J)** Representative HDA-1::GFP images after the L2 symmetric (I) and L2 asymmetric divisions (J). **(K)** Quantification of HDA-1::GFP fluorescence intensity following the L2 symmetric and L2 asymmetric divisions in V1-V4 lineages. Following symmetric division anterior cells (pa) show similar expression of HDA-1 compared to posterior cells (pp) (n = 71 cells per condition). After L2 asymmetric division posterior daughter cells (pap, ppp) show higher HDA-1 expression compared to anterior cells (paa, ppa) (p<0.001 with a two-tailed t-test, n = 61 cells per condition). Seam cell nuclei in C, E and F are labelled using *SCM::GFP*. In H, I and K the membrane of the seam cells is labelled using the apical junction marker *ajm-1p::ajm- 1::mCherry*. Scale bars are 5μm in C, E, F, H, I and J. Error bars in B, D, G and K show the mean ± standard deviation and **** represent *p*<0.001, *** *p*<0.005, ** *p*<0.01 with a t-test.

Correct seam cell patterning depends on a divergent version of the canonical Wnt signalling pathway called the Wnt/β-catenin asymmetry pathway (WβA), which ensures that following asymmetric division, posterior daughters activate Wnt target genes and retain seam cell fate, while anterior daughters repress Wnt targets and differentiate (Jackson & Eisenmann, 2012; Sawa & Korswagen, 2013). Multiple Wnt ligands, including LIN-44, CWN-1, CWN-2, EGL-20, and MOM-2, are expressed in distinct body regions and act redundantly to control the orientation of seam cell polarity along the anterior-posterior axis (Harterink et al., 2011; Pani & Goldstein, 2018; Sawa et al., 2026; Yamamoto et al., 2011). Wnt receptors, including the Frizzled receptors LIN-17 and MOM-5 and the Ror receptor CAM-1, are essential for establishing cellular polarity (Forrester et al., 1999; Goldstein et al., 2006; Sawa et al., 1996, 2026; Yamamoto et al., 2011). In dividing seam cells, Wnt signalling promotes cortical localisation of Wnt receptors and Dishevelled proteins, while negative regulators such as PRY-1/Axin and APR-1/APC are enriched at the anterior cortex (Baldwin & Phillips, 2014; Mizumoto & Sawa, 2007a, 2007b). This asymmetry generates daughter cells with distinct nuclear levels of POP-1/TCF and SYS-1/β-catenin (Mizumoto & Sawa, 2007b). Low POP-1 and high SYS-1 in posterior daughters activate Wnt target genes, whereas high POP-1 and low SYS-1 in anterior daughters lead to transcriptional repression (Bekas & Philips, 2022; Huang et al., 2007; Vora & Phillips, 2015).

While transcriptional regulation of seam cell development has been extensively studied, the role of chromatin-based mechanisms in this process remains largely unexplored. Components of the SWI chromatin remodelling complex have been implicated in asymmetric division of the T cell lineage (Cui & Han, 2007; Shibata et al., 2012), and the acetylated-histone-binding protein BET-1 contributes to fate maintenance in T cell progeny (Shibata et al., 2010). However, the broader impact of chromatin modifications on the expression of seam cell regulators remains unclear. Class I histone deacetylases (HDACs) regulate gene expression by removing acetyl groups from histone tails, promoting chromatin compaction and transcriptional repression (Seto & Yoshida, 2014; Shahbazian & Grunstein, 2007). These enzymes typically function within multiprotein repressor complexes such as SIN3, NuRD, and CoREST, and play crucial roles in promoting cellular differentiation during animal development (Ahringer, 2000; Brunmeir et al., 2009; Dovey et al., 2010b; Jaju Bhattad et al., 2020). The *C. elegans* class I histone deacetylase HDA-1, homologous to mammalian HDAC1/2, has been implicated in the regulation of multiple developmental processes. During embryogenesis, HDA-1 acts together with UNC-37/Groucho as a POP-1 co-repressor downstream of the Wnt/β-catenin asymmetry pathway to repress endoderm fate in the MS blastomere (Bekas & Philips, 2022; Calvo et al., 2001). HDA- 1 also contributes to nervous system development by regulating cell migration and axon guidance (Zinovyeva et al., 2006) and, as part of the SIN3 complex, participates in the formation of male sensory rays (Choy et al., 2007). Furthermore, HDA-1 plays critical roles in gonadogenesis and vulval development, including promoting the invasive behaviour of the anchor cell (Matus et al., 2015), and preventing ectopic vulval fate specification as part of the SynMuvB pathway (Dufourcq et al., 2002; Solari & Ahringer, 2000). Transcriptional profiling in the epidermis has shown that *hda-1* is expressed in seam cells and that its downregulation affects terminal seam cell number (Katsanos et al., 2021). Despite these observations, the specific role of HDA-1 in regulating stem cell behaviour and asymmetric division has not been explored.

In this study, we identify a role for HDA-1 in regulating seam cell polarity orientation in *C. elegans*. We report that loss of *hda-1* disrupts the orientation of asymmetric divisions and results in defects in molecular asymmetry between daughter cells. Using Targeted DamID we identify the Wnt receptors *lin-17*/Frizzled and *cam- 1*/Ror as HDA-1 targets. Single-molecule FISH (smFISH) analysis reveals elevated expression of *lin-17* and *cam-1* in *hda-1* mutants, disrupting the graded expression pattern observed in wild type and reducing the differential expression between anterior and posterior daughters. Furthermore, we show that overexpression of these Wnt receptors is sufficient to phenocopy the *hda-1* mutant polarity defects. Together, our findings uncover a novel role for HDA-1 in ensuring proper orientation of asymmetric divisions through regulation of Wnt receptor expression.

## Results

### *hda-1* expression is largely uniform in seam cells

We previously identified through transcriptional profiling *hda-1* as a seam cell- expressed chromatin regulator. We reported that RNAi-mediated downregulation of *hda-1* using a seam cell-driven hairpin construct caused seam cell hyperplasia (Katsanos et al., 2021). This phenotype is specific to *hda-1* as RNAi targeting of other HDAC family members did not lead to statistically significant changes in seam cell number (Figure 1B), so we decided to characterise in more detail.

To investigate the expression pattern of *hda-1* in different seam cells, we first quantified *hda-1* transcripts using smFISH. *hda-1* mRNA was detected in all seam cell lineages and surrounding hypodermal cells (Figure 1C, E-F). At the late L1 stage, transcript abundance was highest in the mid-body V1-V5 seam cells, with lower levels in the H and T lineages at the head and tail (Figure 1C-D). We next examined *hda-1* expression during V1-V4 seam cell divisions. Following the L2 symmetric division, *hda-1* transcript levels were similar between anterior and posterior daughters (Figure 1E, G). After the L2 asymmetric division, *hda-1* mRNA remained detectable in both daughters, with only mild and variable enrichment in anterior cells (Figure 1F-G), hence *hda-1* transcript is largely uniform between seam cell daughters post division.

To assess HDA-1 distribution at the protein level, we analysed an endogenous HDA-1::GFP reporter. HDA-1::GFP localised to the nuclei of all seam cells and hypodermal cells (Figure 1H) with levels that were comparable between daughters after symmetric division (Figure 1I, K). However, after asymmetric division, HDA- 1::GFP showed moderate enrichment in posterior daughter cells (Figure 1J, K). This pattern is not reflected at the mRNA level so it suggests that HDA-1 may be subject to translational or post-translational regulation, as has been reported in other contexts (Poulin et al., 2005; Sengupta & Seto, 2004). Such regulation could affect its stability, activity or interactions with partners during asymmetric division

### *hda-1* mutants display reversals in the polarity of asymmetric divisions

To investigate the function of HDA-1 in seam cell development, we generated a seam cell-specific *hda-1* mutant using the Cre-lox recombination system. We confirmed that neither the introduction of loxP sites flanking the *hda-1* locus nor the expression of Cre recombinase in the seam cells disrupted seam cell development (Figure S1A). Furthermore, *hda-1*^loxP^ animals expressing Cre recombinase showed complete loss of *hda-1* transcripts in seam cells by smFISH, while expression remained detectable in the surrounding hypodermis, confirming the tissue specificity of the knockout (Figure S1B).

Seam cell-specific *hda-1* mutants exhibited clear patterning defects, with both extra cells and gaps in the seam cell line at the early adult stage, suggesting that loss of *hda-1* disrupts the balance between seam cell self-renewal and differentiation (Figure 2A). Quantification revealed an overall increase in mean seam cell number (Figure 2B). To determine when seam cells are gained or lost in *hda-1* mutant development, we scored seam cell number at late L2, L3 and L4 stages. We found that changes in seam cell number were restricted to the L4 stage (Figure 2B).

**Figure 2.**
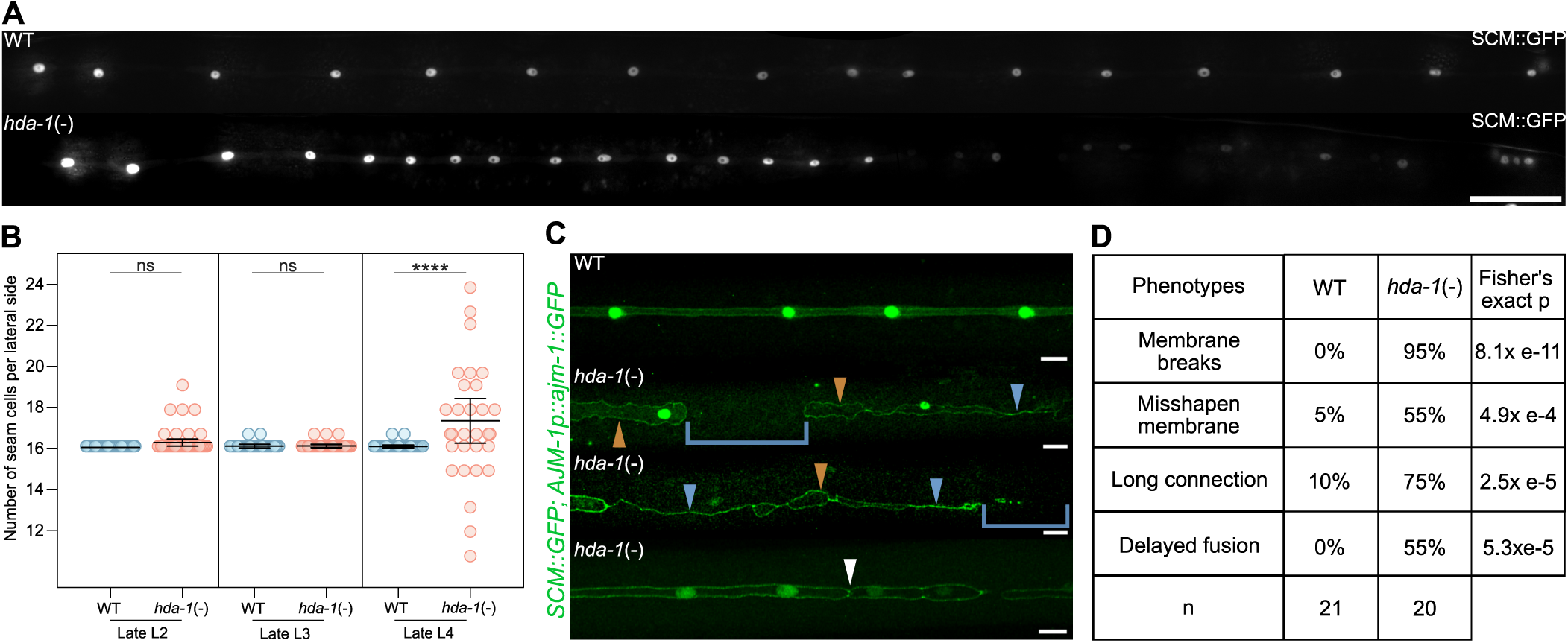
A seam cell-specific *hda-1* mutant displays gains and losses of seam cells. **(A)** Representative images of tissue-specific *hda-1* mutant animals (simplified here as *hda-1*(-)) versus floxed animals not expressing the Cre recombinase (WT) at the early adult stage. **(B)** Seam cell counts in tissue-specific *hda-1* mutant animals at late L2, late L3 and late L4 stages. Only late L4 mutants show a significant increase in the average number of seam cells in comparison to WT (p<0.001 with a two-tailed t-test, n ≥ 58 animals per condition). Error bars show the mean ± standard deviation. **(C)** Representative images of *hda-1* mutant defects at the early adult stage. Membrane breaks are represented with a blue bracket, misshapen membrane with an orange arrow, long connections with a blue arrow and membrane boundaries indicative of delayed cell fusion with a white arrow. The membrane of the seam cells is labelled using the apical junction marker *ajm-1p::ajm-1::GFP*. **(D)** Table showing the frequency of these defects in *hda-1* mutants (n = 20 animals) compared to WT (n = 21 animals). In A and C, seam cell nuclei are labelled using *SCM::GFP*. Scale bars are 50 μm in A and 10 μm in C. See also Figure S1.

At the late L4-early adult stage, seam cells normally fuse to form a syncytium, which can be visualised with the apical junction AJM-1::GFP marker as two perpendicular membrane lines with seam cell nuclei enclosed within (Sulston & Horvitz, 1977). In *hda-1* mutants, however, we observed an array of defects, including membrane breaks, misshapen membrane, long connections between separated cells, and delayed fusion with persistent boundaries between seam cells (Figure 2C). These defects were highly penetrant, with 95% of *hda-1* mutant animals displaying at least one type of abnormality (Figure 2D). These strong phenotypes by the L4 stage were reminiscent of polarity defects and led us to explore whether earlier errors might also be present that do not directly impact seam cell number.

To investigate whether the stereotypic polarity of asymmetric divisions was affected in *hda-1* mutants, we examined expression of the fusogen *eff-1*, which is normally restricted to differentiating anterior daughter cells following the L2 asymmetric division (Koneru et al., 2021). In wild-type animals, *eff-1* expression was exclusively observed in anterior daughter cells, as expected (Figure 3A). In contrast, *hda-1* mutants showed reversed *eff-1* expression in some lineages, with posterior daughters expressing *eff-1* while anterior daughters did not (Figure 3A). Quantification revealed that 18% of *hda-1* mutants displayed this reversal in the V2 lineage (Figure 3B).

**Figure 3.**
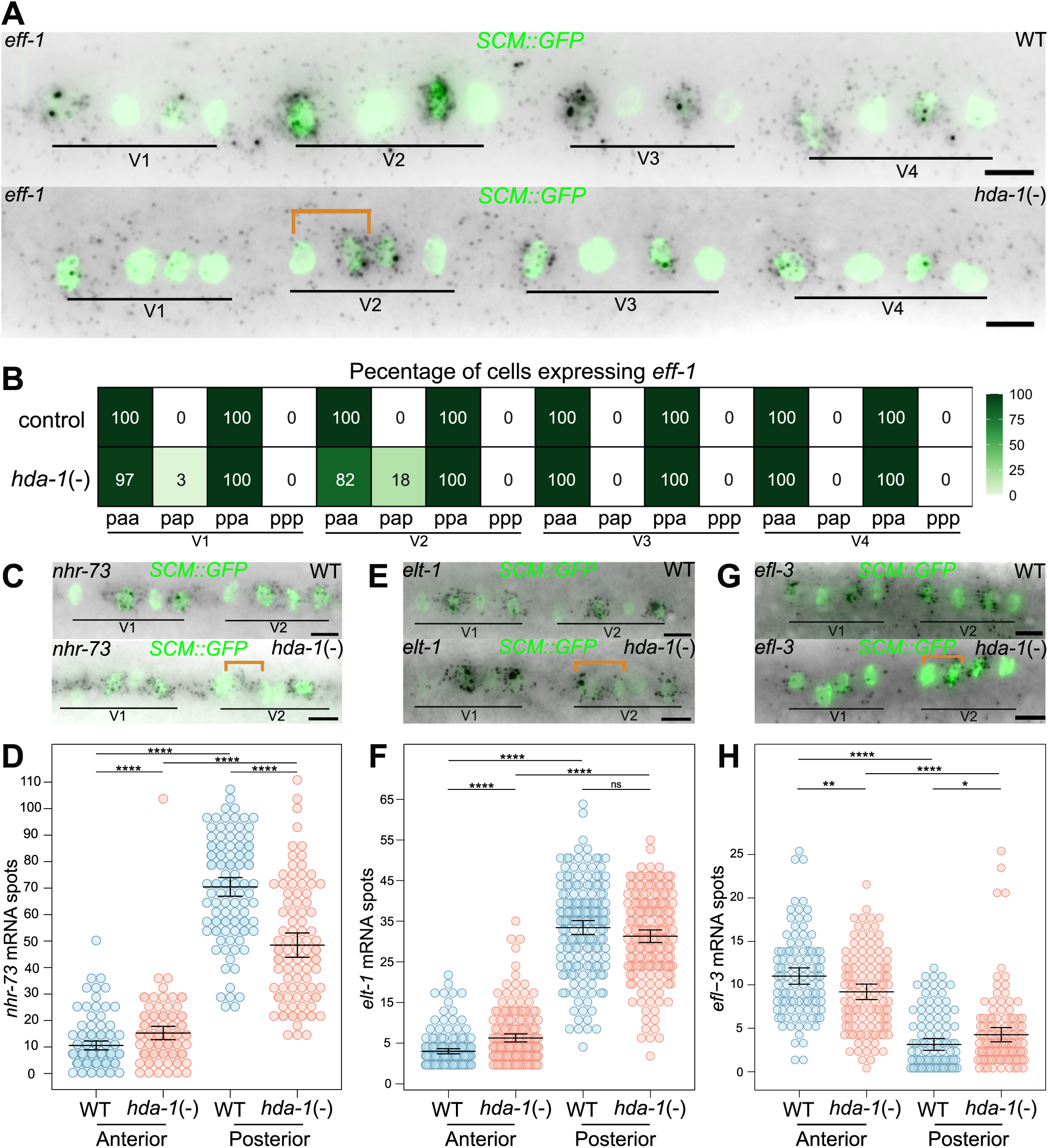
*hda-1* mutants show polarity reversals upon asymmetric cell division. **(A)** Representative images of *eff-1* smFISH following the L2 asymmetric division in WT and in tissue-specific *hda-1* mutant animals (*hda-1*(-)). Orange bracket indicates a division with reversed expression of *eff-1*. **(B)** Heatmap showing frequency of cells expressing *eff-1* following the L2 asymmetric division in WT and *hda-1* mutants (26 ≤ n ≤ 38 cells per condition). **(C-H)** Representative images (C, E, G) and quantification (D, F, G) of mRNA spots in WT (blue) and in *hda-1* mutant animals (red) following the L2 asymmetric division for *nhr-73* (C-D) *elt-1* (E-F) and *efl-3* (G-H). Orange brackets in C, E, G show a pair with reversed expression of the corresponding gene. 94 ≤ n ≤ 110 cells in D, 170 ≤ n ≤ 174 cells in F and 104 ≤ n ≤ 114 cells in G. In A, C, E and G seam cell nuclei are labelled using *SCM::GFP* and scale bars are 5 μm. Error bars in D, F and H show the mean ± standard deviation and **** represent *p*<0.001, ** *p*<0.01, * *p*<0.05 with a two-tailed t-test.

We next investigated whether molecular asymmetry between daughter cells was affected by assessing expression of genes that are differentially expressed between anterior and posterior daughter cells following asymmetric division. We examined *nhr-73* and *elt-1*, which are normally enriched in posterior daughter cells (Miyabayashi et al., 1999; Smith et al., 2005), and *efl-3*, which is enriched in anterior daughter cells (Ferrando-Marco & Barkoulas, 2025) (Figure 3C, E, G). In all cases, we found that molecular asymmetry between daughter cells was significantly reduced in *hda-1* mutants (Figure 3D, F, H). Taken together, these findings suggest that loss of *hda-1* leads to polarity defects and impaired molecular asymmetry during asymmetric division, which precede and likely contribute to the gains and losses of seam cells observed at later developmental stages.

### Identification of putative HDA-1 targets in seam cells via TaDa

To identify downstream targets of HDA-1 in the seam cells, we performed Targeted DamID (TaDa) profiling. We expressed an *hda-1::dam* fusion at very low levels using a bicistronic TaDa cassette under the seam-cell promoter *srf-3i*, following established approaches to minimise Dam-associated toxicity (Katsanos & Barkoulas, 2022; Southall et al., 2013). Indeed, expression of the *hda-1::dam* fusion did not affect developmental speed or brood size (Figure S2A-B). A nuclear GFP-Dam fusion expressed from the same cassette served as the control. Animals were collected at the late L3 stage, and genomic regions methylated by HDA-1::Dam were isolated and sequenced.

Aggregate genome-wide signal profiles across genes containing peaks between 5 kb upstream of the transcriptional start site (TSS) and 2kb downstream of the transcription end site (TES) showed that HDA-1 binding is enriched within gene bodies (Figure S2C). We identified 1,886 statistically significant peaks (FDR < 0.05) (Table S1). Hierarchical clustering of the localisation and score of statistically significant peaks within genes revealed clusters of signal enrichment upstream of TSS, inside genes, and downstream of TES regions for HDA-1 TaDa (Figure S2D). Peaks were assigned to the closest gene leading to 3,196 HDA-1 putative targets. The list of candidate HDA-1 target genes showed significant overlap with genes expressed in seam cells based on single-cell combinatorial indexing RNA-sequencing data (Cao et al., 2017) and with a lab-curated list of genes with known roles in seam cell development from the literature (Table S2). Gene ontology (GO) term analysis of putative HDA-1 targets in the seam cells revealed terms relating to development, Wnt signalling and transcription (Figure S2E).

### Loss of HDA-1 function disturbs the graded expression pattern of the Wnt receptors *lin-17* and *cam-1*

To study how HDA-1 may exert its effect, we focused on Wnt signalling, which has been previously linked to seam cell polarity. We selected several candidates identified through TaDa for expression analysis by smFISH, including the Wnt receptors *lin-17*, *cam-1*, and *mom-5*, as well as the Dishevelled gene *dsh-2* and the negative regulators *apr-1* and *pry-1* (Figure S2F). Among all candidates examined, *lin- 17* and *cam-1* showed the most striking and consistent expression changes in *hda-1* mutants. As the mRNA expression patterns of these Wnt receptors had not been previously quantified in seam cells, we analysed *cam-1* and *lin-17* expression in wild-type animals. First, *cam-1* expression was enriched in anterior seam cell lineages, with the highest expression in V1 and progressively lower levels in V2, V3, and V4 at late L1 and following the L2 symmetric and asymmetric divisions (Figure S3A-D). Interestingly, following symmetric and asymmetric cell divisions, it was the anterior daughter cells that showed significantly higher *cam-1* expression compared to their posterior counterparts (Figure S3D). Analysis of *cam-1* mRNA distribution more broadly across the lateral epidermis was consistent with graded expression along the anterior-posterior axis with highest expression in the head and mid-body regions and lowest expression in the tail (Figure S3E). In *hda-1* mutants, *cam-1* transcript levels were significantly elevated in seam cells following both the L2 symmetric and asymmetric divisions (Figure 4A-D). This upregulation was observed across all Vlineages and in both anterior and posterior daughter cells, indicating that HDA-1 normally restricts *cam-1* expression. As a consequence, *hda-1* mutants showed significantly reduced anterior-posterior (A-P) asymmetry for *cam-1* expression (Figure 4E) and reduced V1-V4 differences compared to wild-type (Figure 4E). This indicates that upregulation of *cam-1* disrupts differences across lineages and asymmetric expression between daughter cells.

**Figure 4.**
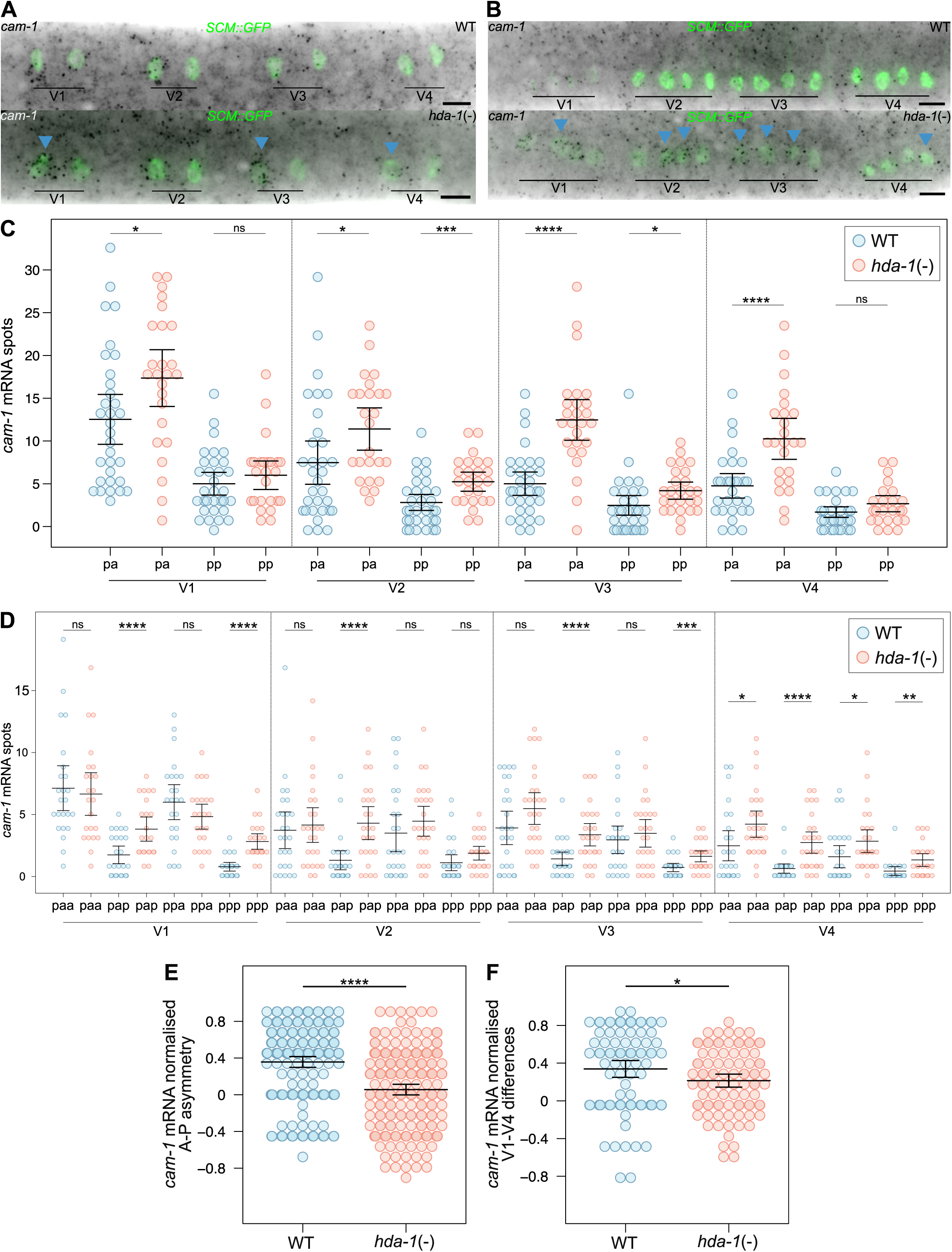
*cam-1* expression is increased in *hda-1* mutants. (A-B) Representative *cam-1* smFISH images of WT and tissue-specific *hda-1* mutant animals (*hda-1*(-)) following the L2 symmetric (A) and L2 asymmetric divisions (B). Blue arrowheads show examples of cells with visually higher *cam-1* mRNA spots. Seam cell nuclei are labelled using *SCM::GFP* and scale bars are 5 μm. **(C-D)** Quantification of *cam-1* mRNA spots in V1-V4 lineages in WT (blue) and *hda-1* mutant animals (red) following the L2 symmetric (C) and L2 asymmetric divisions (D) (in C 24 ≤ n ≤ 32, and in D 24 ≤ n ≤ 27 cells per condition). **(E)** Normalised anterior-posterior (A-P) asymmetry of *cam-1* mRNA following the L2 asymmetric division in WT (blue) and *hda-1* mutant animals (red). A-P asymmetry was calculated for each anterior/posterior sister-cell pair as a normalised difference (A-P)/(A+P+1), where values close to 0 indicate equal expression between sisters, positive values indicate higher expression in anterior cells, and negative values indicate higher expression in posterior cells (202 ≤ n ≤ 206 cell pairs per condition). **(F)** Normalised V1-V4 lineage differences in *cam-1* mRNA following the L2 asymmetric division in WT (blue) and *hda-1* mutants (red). V1-V4 differences were calculated for matched cell types (paa, pap, ppa, ppp) using a normalised difference (V1-V4)/(V1+V4) where values close to 0 indicate equalisation of expression between anterior (V1) and posterior (V4) lineages (n = 92 cell pairs per condition). Error bars in C-F show the mean ± standard deviation. In C and D, **** represent *p*<0.001, *** *p*<0.005, ** *p*<0.001 and * *p*<0.05 with a two-tailed t-test. In E and F, * p<0.05 **** p<0.001 with a Mann-Whitney test. See also Figure S3.

Regarding *lin-17,* WT animals showed enrichment in posterior seam cell lineages, with V4 showing the highest expression and V1 the lowest among V1-V4 following the L2 symmetric and asymmetric divisions (Figure S4A-D). In contrast to what was seen for *cam-1* expression, posterior daughter cells showed significantly higher *lin-17* expression compared to their respective anterior daughters after symmetric and asymmetric divisions (Figure S4D). Analysis of *lin-17* mRNA across the lateral epidermis further confirmed enrichment in the posterior region of the body (Figure S4E). In *hda-1* mutants, *lin-17* expression was significantly increased following both the L2 symmetric and asymmetric divisions (Figure 5A-D). This increase was observed across all V lineages analysed following the L2 symmetric division and mostly in V1 and V2 following the L2 asymmetric division. Analysis of A-P asymmetry revealed that *lin-17* expression differences between anterior and posterior daughters remained largely unchanged in *hda-1* mutants compared to wild type animals (Figure 5E). However, examination of V1-V4 differences showed that *hda-1* mutants displayed significantly reduced differences across lineages compared to wild-type (Figure 5F), indicating that HDA-1 loss disrupts mostly the posterior-anterior gradient of *lin-17* expression while largely preserving daughter cell asymmetry.

**Figure 5.**
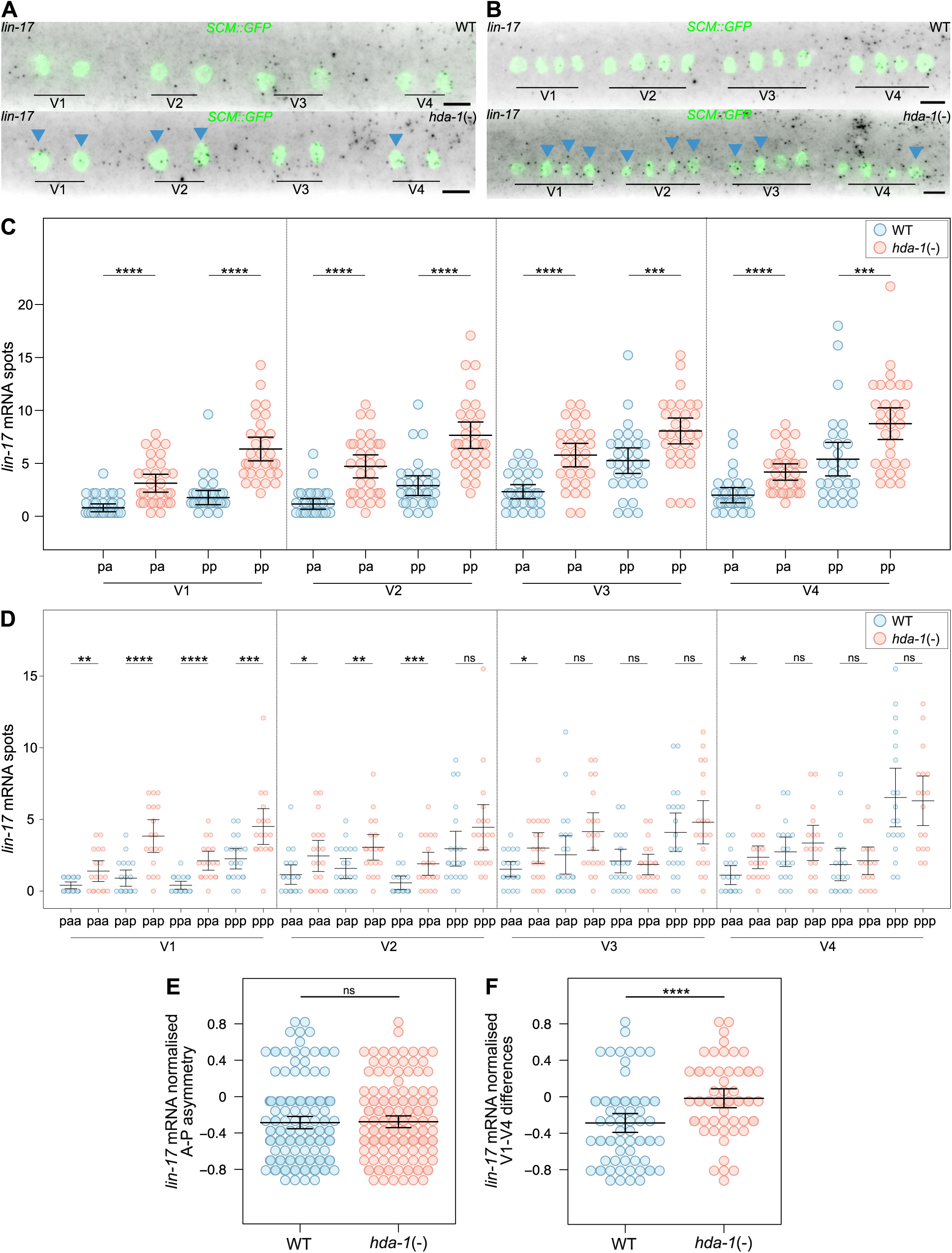
*lin-17* expression is increased in *hda-1* mutants. (A-B) Representative *lin-17* smFISH images of WT and tissue-specific *hda-1* mutant animals (*hda-1*(-)) following the L2 symmetric (A) and L2 asymmetric divisions (B). Blue arrowheads show examples of cells with visually higher *lin-17* mRNA spots. Seam cell nuclei are labelled using *SCM::GFP* and scale bars are 5 μm. **(C-D)** Quantification of *lin-17* mRNA spots in V1-V4 lineages in WT (blue) and *hda-1* mutant animals (red) following the L2 symmetric (C) and L2 asymmetric divisions (D) (in C 30 ≤ n ≤ 32, and in D 17 ≤ n ≤ 21 cells per condition). **(E)** Normalised anterior–posterior (A–P) asymmetry of *lin-17* mRNA following the L2 asymmetric division in WT (blue) and *hda-1* mutant animals (red). A-P asymmetry was calculated as in Figure 4, (150 ≤ n ≤ 162 cell pairs per condition). **(F)** Normalised V1-V4 lineage differences in *lin-17* mRNA following the L2 asymmetric division in WT (blue) and *hda-1* mutants (red). V1-V4 differences were calculated as in Figure 4 (60 ≤ n ≤ 72 cell pairs per condition). Error bars in C-F show the mean ± standard deviation. In C and D, **** represent *p*<0.001, *** *p*<0.005, ** *p*<0.001 and * *p*<0.05 with a two-tailed t-test. In E and F, * p<0.05 **** p<0.001 with a Mann-Whitney test. See also Figure S4.

In contrast, other Wnt signalling pathway components explored showed weaker or no significant changes in transcript abundance in the *hda-1* mutant background. *mom-5* expression was elevated at the late L1 stage in *hda-1* mutants but showed no significant changes following the L2 divisions (Figure S5). *dsh-2* displayed modest decrease in expression in *hda-1* mutants following the L2 symmetric and asymmetric divisions (Figure S6), while *apr-1* and *pry-1* expression levels showed both a modest decrease following the L2 symmetric division that was not consistent across lineages (Figure S7-S8). Together, these findings indicate that HDA-1 influences expression of multiple Wnt pathway genes, with *cam-1* and *lin-17* showing the most robust and consistent changes in *hda-1* mutants.

### Overexpression of *lin-17* and *cam-1* in seam cells phenocopies the *hda-1* mutant polarity defects

The upregulation of *lin-17* and *cam-1* expression in *hda-1* mutants suggested that increased Wnt receptor expression may underlie the polarity defects observed in these animals. To test this hypothesis, we decided to overexpress *cam-1* and *lin-17* under the *arf-5i* element, which is contained within the SCM enhancer and is sufficient to drive uniform expression in seam cells (Ashley et al., 2021; Katsanos et al., 2021). Overexpression of *lin-17* alone led to a significant increase in seam cell number at late L4, while overexpression of *cam-1* alone did not significantly alter the average seam cell number but increased phenotypic variability (Figure 6B). Co-overexpression of both *lin-17* and *cam-1* resulted in a pronounced increase in seam cell number, with animals simultaneously displaying gaps in the seam cell line (Figure 6A-B). Phenotypic analysis across developmental stages revealed that, similarly to *hda-1* mutants, changes in seam cell number predominantly occurred at the L4 stage (Figure 6B).

**Figure 6.**
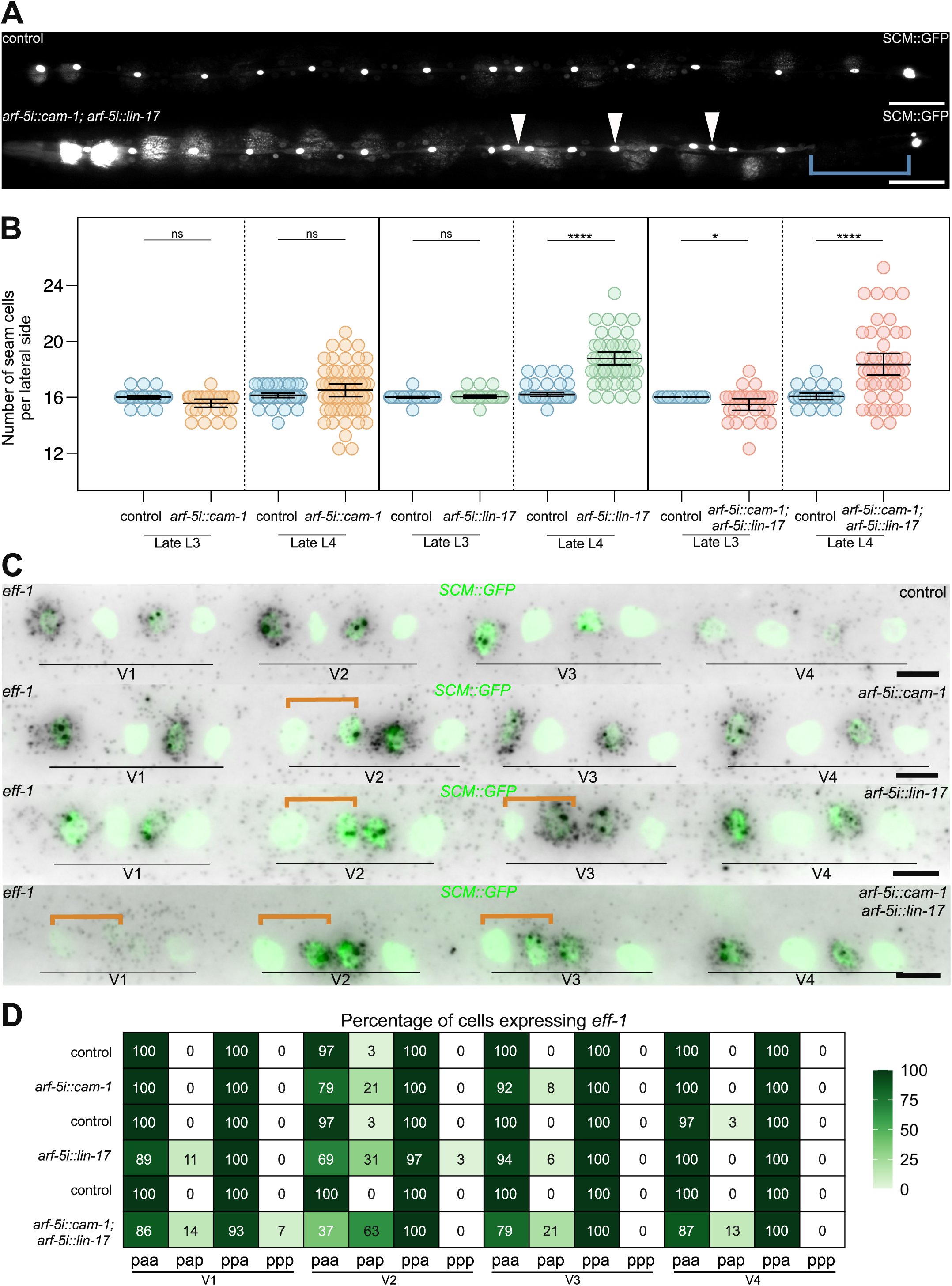
Overexpression of *cam-1* and *lin-17* is sufficient to cause polarity defects. **(A)** Representative images of control and animals overexpressing *cam-1* and *lin-17* under the *arf-5i* element (*arf-5i::cam-1; arf-5i::lin-17*). Blue brackets indicate gaps and white arrowheads extra seam cells. **(B)** Seam cell counts in wild-type (control) and animals overexpressing *cam-1* (orange), *lin-17* (green) and both *cam-1* and *lin-17* (red) under the *arf-5i* element at the late L3 and late L4 stages. Error bars show the mean ± standard deviation and * *p*<0.05 and **** *p*<0.001 with a two-tailed t-test, 30 ≤ n ≤ 60 animals per condition. **(C)** Representative images of *eff-1* smFISH following the L2 asymmetric division in WT and in control and animals overexpressing *cam-1*, *lin-17* and both *cam-1* and *lin-17*. Orange bracket indicates a division with reversed expression of *eff-1*. **(D)** Heatmap showing frequency of cells expressing *eff- 1* following the L2 asymmetric division in the conditions in C (14 ≤ n ≤ 37 cells per condition). In A and C, seam cell nuclei are labelled using *SCM::GFP* and scale bars are 50 μm in A and 5 μm in C.

To assess whether overexpression of these Wnt receptors recapitulates the polarity defects observed in *hda-1* mutants, we examined *eff-1* expression following the L2 asymmetric division. Overexpression of either *lin-17* or *cam-1* individually resulted in polarity reversals closely resembling those of *hda-1* mutants, with *eff-1* expression detected in posterior daughters with no expression in anterior daughters (Figure 6C). Such reversals were rarely observed in control animals (0-3% frequency). Notably, the V2 lineage, which was most prominently affected in *hda-1* mutants, showed the highest frequency of polarity reversals upon Wnt receptor overexpression, with 21% of animals displaying anterior V2 reversals for *cam-1* overexpression and 31% for *lin-17* overexpression (Figure 6D). Co-overexpression of both receptors further increased the frequency of V2 reversals to 63% and additionally induced polarity defects in other seam cell lineages, although V2 remained the most frequently affected (Figure 6C-D). Taken together, these results show that increasing the expression of Wnt receptors in seam cells is sufficient to phenocopy the polarity defects observed in *hda-1* mutants.

### NuRD and SIN3 disruption does not fully phenocopy *hda-1* polarity defects

HDA-1 is known to function as part of the NuRD and SIN3 chromatin-remodelling complexes (Ahringer, 2000). To investigate whether these complexes mediate the HDA-1 role in polarity regulation, we first examined seam cell phenotypes in mutants of key complex components. Loss of the NuRD components *dcp-66* and *egl-27* resulted in an increase in the average seam cell number (Figure 7A). By contrast, mutants of the NuRD component *lin-53* and the SIN3 component *sin-3* did not show significant differences compared to wild type (Figure 7A). To determine whether HDA-1 may work as part of the NuRD or SIN3 complexes to control polarity, we assessed *eff-1* expression following the L2 asymmetric division in these mutants. Neither *egl-27*, *dcp-66*, nor *sin-3* mutants displayed polarity reversals (Figure 7B-C) similar to *hda-1* mutants (Figure 3B). This suggests that while these chromatin-remodelling complexes play a role in seam cell patterning, they may not mediate the polarity defects caused by loss of HDA-1.

**Figure 7.**
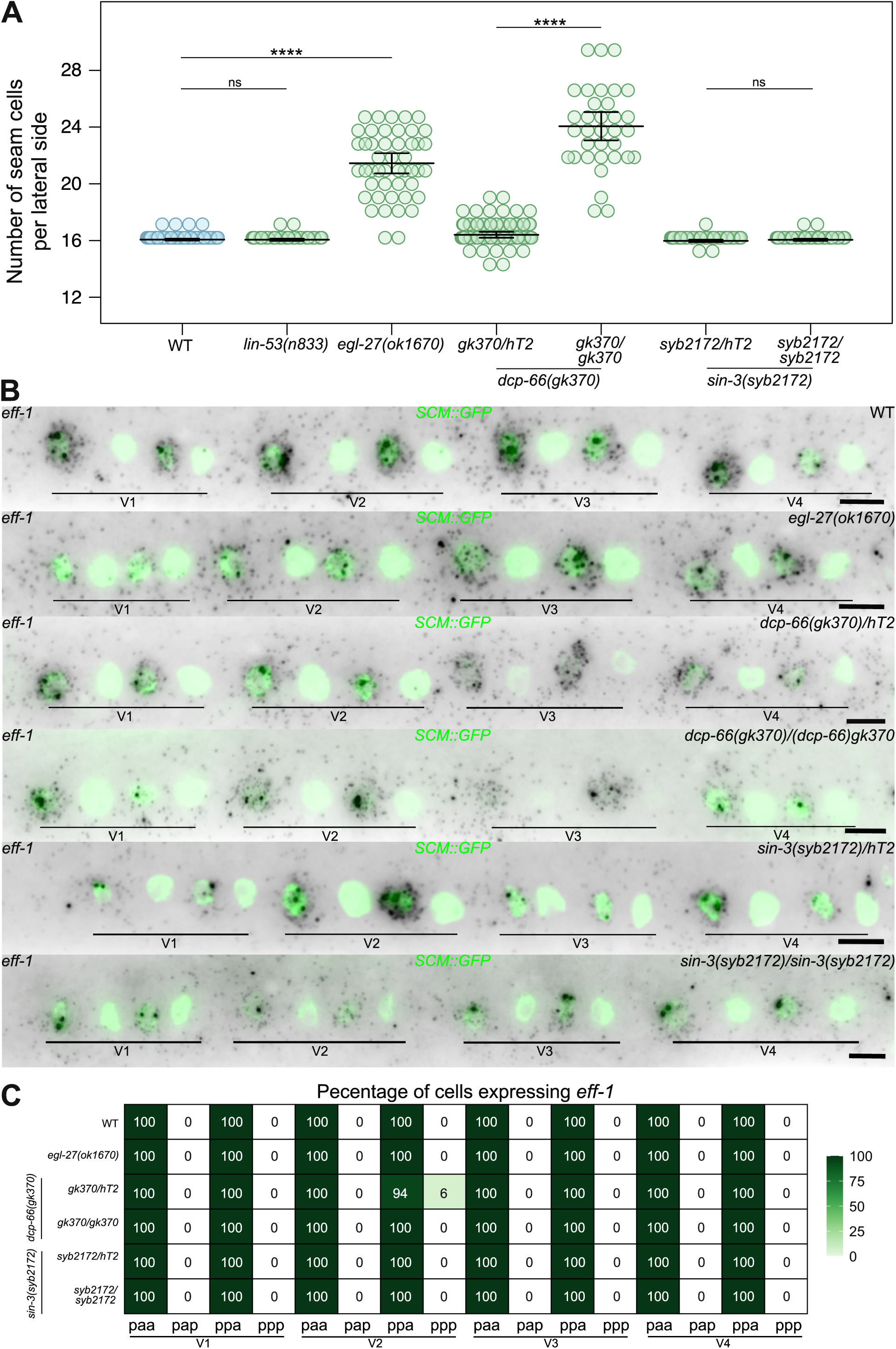
NuRD and SIN3 disruption does not phenocopy the polarity defects of. *hda-1* **mutants. (A)** Seam cell counts in wild-type, and in *lin-53(n833)*, *egl-27(ok1670)*, *dcp-66(gk370)* and *sin-3(syb2172)* mutants at the late L4 stage. Error bars show the mean ± standard deviation and **** *p*<0.001 with a two-tailed t-test, 34 ≤ n ≤ 76 animals per condition. **(B)** Representative images of *eff-1* smFISH following the L2 asymmetric division in WT and in *egl-27(ok1670)*, *dcp-66(gk370)* and *sin-3(syb2172)* mutants. Seam cell nuclei are labelled using *SCM::GFP* and scale bars are 5 μm. **(C)** Heatmap showing frequency of cells expressing *eff-1* following the L2 asymmetric division in the conditions in B (9 ≤ n ≤ 28 cells per condition).

## Discussion

Histone deacetylases are well-established regulators of cell fate specification (Brunmeir et al., 2009; Dovey et al., 2010a; Jaju Bhattad et al., 2020; Jeong-Heon et al., 2004). Previous work on *hda-1* in *C. elegans* has shown roles in endoderm specification, development of the nervous system, sex-specific gene expression and vulval patterning (Calvo et al., 2001; Dufourcq et al., 2002; Grote & Conradt, 2006; Matus et al., 2015; Ranawade et al., 2013; Zinovyeva et al., 2006). We report here that HDA-1 regulates asymmetric cell division polarity in the *C. elegans* epidermal seam cells by repressing the graded expression of the Wnt receptors *lin-17* and *cam-1*. In seam cells, Wnt receptors have been shown to be redundantly required for establishment of polarity, as simultaneous disruption of *cam-1*, *lin-17* and *mom-5* results in significant loss of polarity (Yamamoto et al., 2011). More recently, CAM-1 and LIN-17 have been reported to play crucial roles in polarity orientation during seam cell divisions although how they contribute to this process remains unclear (Sawa et al., 2026). Our study therefore reveals a link between chromatin-mediated transcriptional repression and Wnt-regulated polarity orientation in the context of asymmetric cell division.

Our findings demonstrate that *cam-1* and *lin-17* display graded expression patterns along the anterior-posterior axis. While *cam-1* exhibits an anterior-to-posterior decreasing gradient, *lin-17* shows an opposite posterior-to-anterior gradient, whereas *mom-5* appears to be expressed more in the mid-body region. Gradients of Wnt receptors have been reported in other systems too, for example across the embryonic telencephalon in mice or in the developing wing epithelium in flies, and they are thought to influence graded competence of cells to Wnt ligands or shape the range of Wnt ligand action by limiting diffusion or protecting Wnt ligands from degradation (Cardigan et al., 1998; Chaudhary et al., 2019; Kim et al., 2001). It remains currently unknown how the *cam-1* and *lin-17* gradients are established. HDA-1 shows a relatively uniform expression across seam cell lineages, with enrichment in midbody lineages (V1-V5). This suggests that if HDA-1 contributes to shaping these gradients, it likely does so through interactions with transcription factors that provide positional information along the anterior-posterior axis. Hox genes, which specify positional identity along the body axis and regulate cell fate in a spatially restricted manner, are strong candidates for this role (Mallo & Alonso, 2013). In mammalian cells, Hox proteins have been shown to recruit HDACs through interactions with PBC cofactors to repress transcription (Li et al., 2026; Saleh et al., 2000). In *C. elegans*, several Hox genes show distinct expression domains along the anterior-posterior axis modulating seam cell development (Chisholm & Hsiao, 2012). Notably, Hox genes *lin-39*, *mab-5*, and *ceh-13* are among the putative HDA-1 targets identified in our TaDa analysis. Therefore, HDA-1 may also influence Wnt receptor gradients indirectly through regulation of Hox gene expression. Interestingly, Wnt signalling itself can regulate Wnt receptor gradient establishment through feedback as shown for example in the *Drosophila* wing where Wnt signalling represses *frizzled 2* (*Dfz2*) expression (Cardigan et al., 1998; Sato et al., 1999). This is consistent with the expression patterns observed in daughter cells following L2 division where *cam-1* was found to be enriched in anterior daughters and *lin-17* in posterior daughters. These patterns may reflect negative and positive feedback respectively via Wnt signalling pathway activation.

In *hda-1* mutants, both *cam-1* and *lin-17* receptors are upregulated, disrupting their normal spatial gradients. This disruption correlates with defects in polarity orientation. Consistent with this notion, increased expression of the Wnt receptors *cam-1* and *lin- 17* is sufficient to lead to polarity reversals, supporting a role for spatially patterned receptor expression in polarity orientation control. Recently, overexpression of *lin-17* in a *cam-1;mom-5* mutant background was shown to result in random polarity orientation in V1–V4 seam cell lineages (Sawa et al., 2026), which was taken to suggest that LIN-17 does not itself encode polarity cues. In contrast, our observation that uniform *lin-17* overexpression in an otherwise wild-type background leads to polarity reversals suggests that polarity defects arise from loss of spatial information encoded by receptor distribution. In addition, it has been proposed that Wnt ligands can exert both instructive and permissive effects on polarity orientation, and that the permissive function of Wnt signalling requires additional, unidentified polarity cues (Sawa et al., 2026; Yamamoto et al., 2011). Opposing anterior-posterior gradients of *cam-1* and *lin-17* could provide such intrinsic positional cues that allow seam cells to orient polarity in response to Wnt signals, with the relative levels of different receptors determining how individual cells interpret this signal (Figure 8A). Differential receptor expression between anterior and posterior daughters following division may further refine polarity orientation at the level of individual lineages (Figure 8A). Interestingly, both levels of spatial organisation are disrupted in *hda-1* mutants: differences in receptor expression along the anterior-posterior axis are reduced, and asymmetry between daughter cells is decreased, particularly for *cam-1* (Figure 8B). Wnt receptors are thought to localise to the posterior cortex of the dividing seam cell, leading to asymmetric segregation of downstream regulators such that anterior daughters repress Wnt target genes while posterior daughters activate them (Goldstein et al., 2006; Mizumoto & Sawa, 2007b). An increase in receptor abundance could interfere with this process by destabilising receptor localisation and thereby impairing polarity orientation. However, we note that absolute receptor levels alone cannot fully account for the localisation of polarity defects observed in *hda-1* mutants. If polarity reversals were solely caused by receptor levels exceeding a threshold that disrupts localisation, reversals would be expected to occur most frequently in lineages with the most extreme changes from baseline expression (V1 and V4). However, we found that polarity reversals occur preferentially in V2 across both *hda-1* mutants and receptor overexpression conditions, suggesting that additional factors are likely to influence seam cell sensitivity to polarity defects.

**Figure 8.**
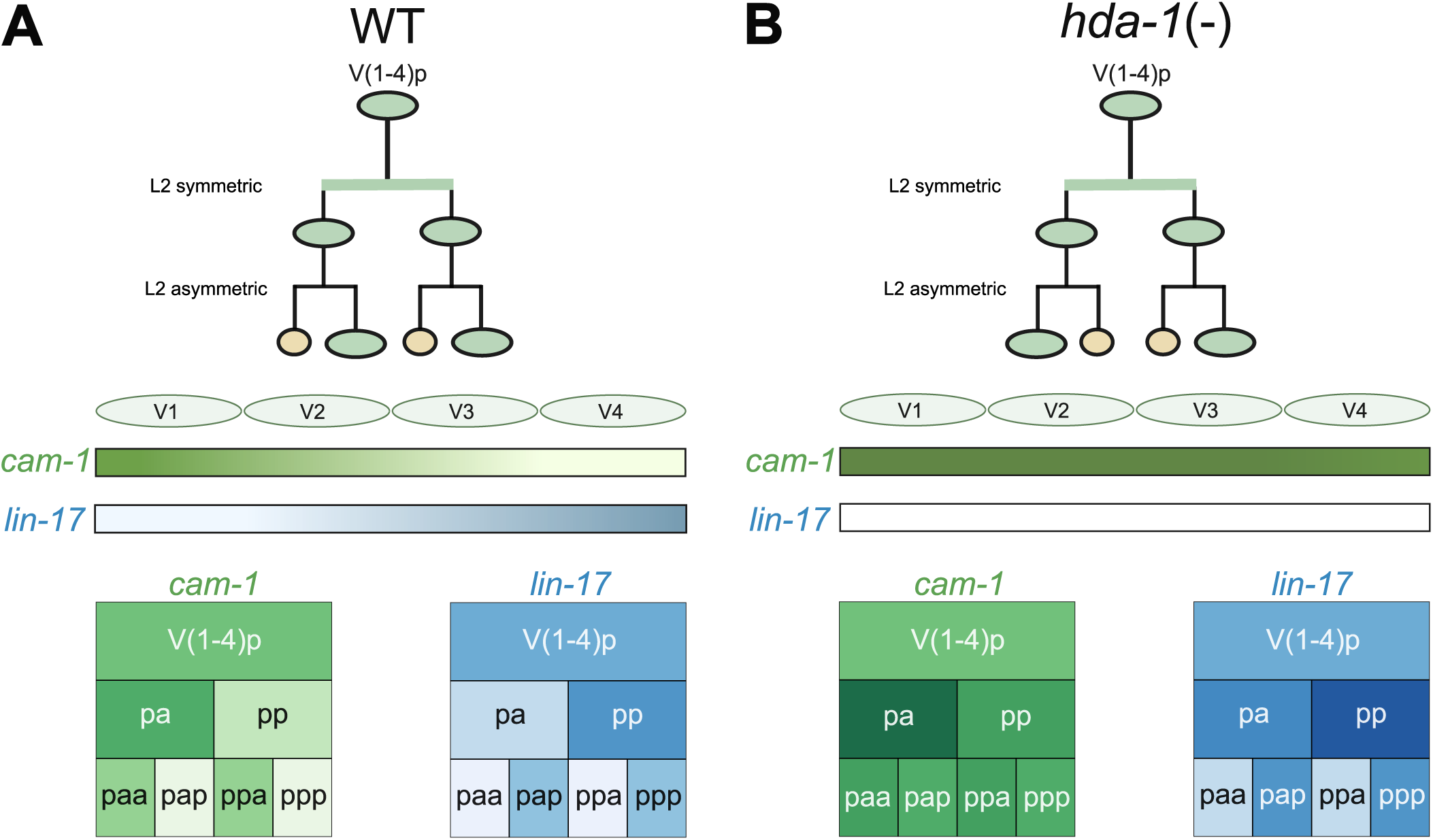
Model for HDA-1 regulation of Wnt receptor expression and polarity orientation in seam cells. **(A)** In wild-type animals, seam cell polarity is consistently oriented along the anterior-posterior axis. The Wnt receptors *cam-1* (green) and *lin-17* (blue) display complementary, graded expression across V1-V4 seam cell lineages, with *cam-1* enriched anteriorly and *lin-17* enriched posteriorly. In addition, within individual lineages, *cam-1* and *lin-17* show differential expression between anterior and posterior daughter cells following division. **(B)** In *hda-1* mutants, asymmetric divisions show polarity reversals and expression of both *cam-1* (green) and *lin-17* (blue) is increased. Differences in receptor expression along the anterior-posterior axis and asymmetry between anterior and posterior daughter cells are reduced. Disruption of graded and asymmetric receptor expression may impair the interpretation of permissive Wnt signals, potentially contirbuting to defects in polarity orientation and polarity reversals.

It is of note that *hda-1* mutants not only displayed polarity reversals without change to the seam cell number at the L2 stage, but also exhibited both extra seam cells and loss of seam cells manifested at late L4. Wnt receptor overexpression was found to be sufficient to phenocopy both early reversals and late stage symmetrisations, raising the possibility that polarity defects can manifest differently depending on the developmental stage. Symmetrisations of asymmetric divisions are often associated with impaired POP-1-mediated repression (Bekas & Philips, 2022; Ferrando-Marco & Barkoulas, 2025). In the embryonic MS blastomere, HDA-1 functions together with UNC-37/Groucho as a co-repressor of the TCF factor POP-1 (Calvo et al., 2001). Given that UNC-37 similarly acts as a POP-1 co-repressor in seam cells (Bekas & Philips, 2022), HDA-1/UNC-37 may also contribute to the repression of Wnt target genes in anterior differentiating daughters through regulation of POP-1. HDACs lack DNA-binding domains and require interactions with DNA- binding proteins or multiprotein complexes to be recruited to specific genomic locations. In *C. elegans*, HDA-1 typically functions as part of the NuRD and SIN3 chromatin-remodelling complexes (Ahringer, 2000; Chen & Han, 2001; Choy et al., 2007). While mutants of NuRD and SIN3 components show changes in terminal seam cell number, they do not display the same L2 polarity reversals. This suggests that, while HDA-1 may cooperate with POP-1 and function as part of the NuRD or SIN3 complex in certain aspects of seam cell gene regulation, its role in establishing correct polarity orientation likely depends on additional interacting partners that remain to be identified.

## Materials and Methods

### *C. elegans* maintenance and genetics

All *C. elegans* strains in this study were maintained at 20°C on Nematode Growth Medium (NGM) plates seeded with *E. coli* OP50 (Brenner, 1974). A list of strains used in this study is provided in Table S3.

### Molecular cloning and transient transgenesis

To overexpress *cam-1* using the last intron of *arf-5* (*arf-5i*), *cam-1^a^* fragment was amplified from N2 cDNA using the oligos cam-1CDS(pes-10) Fw and cam- 1CDS(unc-54UTR) Rv. The amplicon was inserted by Gibson Assembly into pIR5 to create pMFM32. pMFM32 was injected into JR667 at 50 ng/μl with 5 ng/μL *myo- 2::dsRed* as co-injection marker and 55 ng/μl BJ36 as carrier DNA. pMFM32 and pDK5 were injected into JR667 at 40 ng/μl each, with 5 ng/μL *myo-2::dsRed* as co-injection marker and 25 ng/μl BJ36 as carrier DNA.

The TaDa constructs for *hda-1* were built using pDK7. *NLS-GFP* was amplified from pDK54 with NLS:GFPu-DamF and NLS:GFPu-DamR, and *hda-1* was amplified from the fosmid WRM0627aC04 with (mCherry)_hda-1_Fw and (Dam)_hda-1_R. pDK7 was digested with XmaJI and *NLS-GFP* and *hda-1* fragments were inserted upstream and in-frame with Dam to produce pMFM4 and pMFM6. The promoter *srf-3i1::*Δ*pes-*

*10* was introduced using the Multisite Gateway® Technology (Invitrogen). *srf- 3i1::*Δ*pes-10* fragment was amplified from pDK82 using srf-3i_attB4 and delta-pes- 10_attB1r_R. A donor vector was produced via a BP reaction, and was inserted in LR reactions with pMFM4 and pMFM6 to produce 2 different vectors: pMFM8 and pMFM10. A complete list of oligos and vectors used can be found in Table S3 and are available upon request.

### Stable transgenesis

Single-copy transgenic lines were generated by the MosSCI method using animals of the EG6699 strain with a Mos1 transposon insertion on chromosome II (*ttTi5605* locus) showing the uncoordinated (Unc) phenotype (Frøkjær-Jensen et al., 2014). pMFM8 and pMFM10 vectors were injected at 50 ng/μL together with 50 ng/μL of pCFJ601 (Mos1 transposase), 10 ng/μL of pMA122 (heat-shock-inducible *peel-1* toxin) and 10 ng/μl, 2.5 ng/μl and 5 ng/μl of the co-injection markers (pGH8, *myo- 2::dsRed*, *myo-3::mCherry*). After injection animals were kept at 25°C until they were starved. Heat shock was performed at 33°C for 3.5 hours and the plates were allowed to recover for 3 hours at room temperature before ‘reverse chunking’. The next day the top of the laws on each chunk was screened for normal roaming animals (*unc-119*(-) rescued animals) with the absence of co-injection markers. Homozygous lines were confirmed molecularly for the presence of single-copy transgenes using oligos NM3880 and NM3884.

### CRISPR genome editing

*LoxP* sites flanking the *hda-1* endogenous locus (394 bp upstream of the transcription start site and 435 bp downstream of the transcription termination site) were introduced by injecting JR667 animals with 0.25 ng/μL protein Cas-9, 2.25 μM tracrRNA, 2.38 μM of 5’_hda-1_crRNA and 3’_hda-1_crRNAs, 5.5 μM of 5’ and 3’ repair templates and 5 ng/μL *myo-2::dsRed*. crRNAs targeted 414 bp upstream the TSS and 402 bp downstream the TES. 5’ and 3’ *hda-1* repair templates presented 50 bp homology to either side of the Cas-9 cut site and included a *loxP* sequence, a ClaI cut site and a M13uni_43 or M13rev_49 sequence, respectively. Transgenic animals with red pharynx were genotyped using the oligos hda-1_5p_F_2, R and hda-1_3p_F, R for 5’ and 3’ modification, respectively. The PCR product was digested with ClaI restriction enzyme to confirm successful homologous repair.

### RNAi feeding

RNAi by feeding was performed using *E. coli* HT115 expressing double-stranded RNA corresponding to the target gene or the control dsRNA plasmid (empty vector) as food source (Kamath et al., 2003). Bacteria cultures were grown overnight and 300 μl were seeded onto NGM plates containing 25 μg/ml ampicillin, 12.5 μg/ml tetracyclin and 1 mM IPTG. All RNAi treatments were performed during the postembryonic development by seeding eggs from the corresponding strain onto the RNAi plates. RNAi bacteria clones used in this study are from the commercially available Ahringer RNAi Library (Kamath & Ahringer, 2003) (Source Bioscience).

### Analysis of brood size

To measure brood size, animals were synchronised by spot bleaching gravid adults. 48 hours later (day 1), individual L4 stage hermaphrodites were transferred onto 35 mm petri dishes seeded with 100 μl bacterial lawn. 10 individual hermaphrodites were transferred per condition. Animals were allowed to develop into adults and lay self-progeny for 24 hours (I plates). On day 2, animals were transferred onto new 35 mm plates and allowed to lay eggs for another 24 hours (II plates). On day 3, animals were transferred once more to III plates. The number of progeny was scored on subsequent days: plates I at day 3, plates II at day 4 and plates III at day 5. To facilitate scoring, a grid pattern was drawn one the plates using a fine marker (Kwah & Jaramillo-Lambert, 2023). Brood size was calculated by summing the number of progeny produced by each parental. Brood size bar plots represent the mean number of progeny per condition and error bars represent the standard deviation.

### Analysis of developmental time

To measure the developmental time, animals were synchronised by spot bleaching gravid adults. Then, synchronised day 2 adults were let to lay progeny into new NGM plates for 3 hours and then they were removed from the plates. 15 adults were used per plate and three plates representing three biological replicates per condition were set up in parallel. Plates were scored at 45 and 50 hours after removal of the adult animals to find animals that had reached late L4 stage. To identify late L4 animals, the shape of the vulva was used as reference. Bar plots showing developmental time represent average percentage of animals that have reached L4 stage across the three replicates and error bars represent standard error of proportion. To calculate the standard error of proportion, average proportion across replicates (p) and total number of animals scored across replicates (n) was used as follows:

Standard error of proportion: 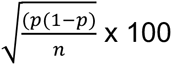

### Phenotypic analysis and microscopy

For seam cell scoring, animals were anesthetised on 3% agarose pads containing 100 μM sodium azide (NaN3), secured with coverslip and visualised under an AxioScope A1 (Zeiss) upright epifluorescence microscope with a LED light source with a RETIGA R6 camera controlled by the Ocular software. The seam cell number was scored on the lateral side closest to the objective.

Protein reporter and membrane marker images were acquired using a Leica SP8 Inverted confocal microscope with a x63/1.4 oil DIC objective controlled by LASX software. Animals were mounted in 5% agarose pads containing 100 μM sodium azide (NaN3) and secured with a coverslip. To visualise L2 symmetric and asymmetric divisions animals were observed at 26 hours post-bleaching. HDA-1::GFP images were acquired in a single plane that provided the best focus for the seam cell nuclei. z-stack slices of 0.4 μm were acquired for animals containing the AJM-1p::AJM- 1::GFP at late L4 stage, 48 hours post-bleaching.

Fluorescence intensity was quantified using FIJI macros developed by the Imperial College FILM facility. The data was normalised (rescaled) to [0,1] using the formula:

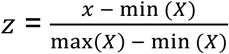

where X is the data, x is a datapoint and z is the corresponding rescaled datapoint. Picture editing was performed using straightening tool and stitching macro from FIJI (Preibisch et al., 2009).

### Single molecule fluorescent in situ hybridisation

Animals were synchronised by bleaching and were fixed at the appropriate stage using 4% formaldehyde for 45 minutes and permeabilised with 70% ethanol. Samples were then incubated with a pool of 24-48 oligos labelled with Quasar 670 (Biosearch Technologies, Novato, CA) for 16 hours. Imaging was performed using the x100 objective of an epifluorescence Ti-eclipse microscope (Nikon) and DU-934 CCD- 17291 camera (Andor Technology) operated via the Nikon NIS Elements Software. 23 z-stack slices of 0.8 μm were acquired for each animal. Spot quantification was performed using MATLAB (MathWorks) as previously described (Raj et al., 2008).

For the epidermal quantification of *cam-1* and *lin-17,* smFISH signal was analysed along the anterior-posterior axis using the epidermal seam cells as anatomical landmarks. Regions of interest (ROIs) were defined between adjacent seam cells (H0-H1, H1-H2, H2-V1, V1-V2, V2-V3, V3-V4, V4-V5 and V5-T). To minimize signal contribution from non-epidermal tissues in the head, mRNA quantification in the two most anterior ROIs (H0-H1 and H1-H2) was restricted to three z-planes corresponding to the epidermal layer, whereas five z-planes were analysed for all more posterior ROIs. ROIs were drawn manually in Fiji, and the area of each ROI was measured in pixel units and converted to physical units using the image calibration (7.6923 pixels = 1 µm). ROI volumes were calculated as the product of ROI area (µm²), z-step size (0.8 µm), and the number of z-planes included for each ROI. For each region, mRNA density was calculated by dividing the number of detected mRNA molecules by the corresponding ROI volume and is reported as mRNAs per 1000 µm³. This volumetric normalisation accounts for differences in ROI size and z-depth across anterior and posterior regions.

For presentation, z-slices containing spots where max-projected to give a single image, inverted and merged to the GFP channel using ImageJ (NIH). For the *eff-1* smFISH experiment in *dcp-66*(-), the seam cell marker signal was adjusted locally in faint nuclei to facilitate lineage identification. A complete list of the smFISH probes used in this study is provided in Table S3.

### DNA extraction and cDNA preparation

Animals from the N2 strain were collected at L2 stage and total RNA was extracted using the TRIzol reagent (Invitrogen) and isopropanol/ethanol precipitation. NanoDrop quantification and gel electrophoresis was used to assess the quantity and quality of RNA. 2 mg of RNA from all samples was subjected to DNase (Promega) treatment and cDNA was synthesized using Superscript IV (Invitrogen) with Oligo(dT) primers as per manufacturer’s instructions.

### Extraction and amplification of methylated DNA

For TaDa experiments animals were synchronised by bleaching and grown for three generations in *E. coli* dam-/- dcm-/- bacteria (New England Biolabs, C2925). Two biological replicates were grown in parallel for strains carrying the TaDa transgenes NLS-GFP::Dam and HDA-1::Dam. Animals were collected after 35 hours post-bleaching, extensively washed x5 in M9 buffer and stored at -20°C. Purification and amplification of methylated DNA was performed as previously described (Katsanos et al., 2021). Extracted gDNA was digested with DpnI and double stranded adaptors were ligated using T4 DNA ligase. Then, DNA fragments were digested with DpnII and amplified using MyTaq polymerase. Amplicons were purified using QIAquick PCR purification kit and adaptors were removed by AlwI digestion followed by another purification. Library preparation and Next Generation Sequencing on an Illumina HiSeq 4000 platform was performed by GENEWIZ.

### NanoDam signal profiles, peak calling, and gene assignment

FASTQ files representing paired-end reads for each sample and replicate were processed using the damidseq_pipeline (v.1.5.3) (Marshall & Brand, 2015). Reads were mapped to the *C. elegans* WBcel235 genome assembly using bowtie2 (v.2.4.5) and scores were calculated per GATC fragment of the genome. Pairwise genome- wide calculations of log2(Dam-fusion/Dam-control) ratio scores per GATC fragment of the genome were performed from bam files to produce bedGraph signal files that were arithmetically averaged into a single signal profile. Peak calling was performed using the perl script find_peaks (available at https://github.com/owenjm/find_peaks) with FDR < 0.05. Statistically significant peaks were assigned to genes using UROPA (Kondili et al., 2017) with Caenorhabditis_elegans.WBcel235.106.gtf as the genome annotation file. Peaks were assigned to genes when their centre coordinate was positioned up to 5 kb upstream of a gene start site or 2 kb downstream of the end site. Visualisation of signal tracks and other genomic features was performed using Integrative Genomics Viewer (IGV).

### Aggregation plots and heatmaps of signal localization

Signal aggregation plots and peak localisation heatmaps were generated using the SeqPlots GUI application (Stempor & Arranger, 2016). Signal and peaks were visualised across genes containing peaks up to 5kb upstream of a gene start site or 2 kb downstream of the end site. Genes were plotted as “anchored” features, where the TSS and the TES are fixed in the x axis and the genic sequence is adjusted to a pseudo-length of 2kb. In aggregation plots, signal is averaged in 10bp bins and y axis represents deviations from the average signal across the plotted region (z-scores). In heatmaps, each coloured line represents the position of a statistically significant peak.

### Gene-set enrichment analysis

Gene-sets identified in this study were assessed for enriched gene ontology (GO) terms using Gene Set Enrichment analysis (WormBase). A *p*-value of <0.005 was used for significance threshold and plots show -log_10_(pvalue).

### Quantification and statistical analysis

Graphic representation and statistical analysis were performed using R. Error bars used in all graphs represent the standard deviation unless otherwise indicated. An unpaired t-test, Mann-Whitney test or chi-square test was used to evaluate significance as indicated in the figure legends. Results were considered statistically significant when p < 0.05. Anterior-posterior asymmetry and V1-V4 differences were quantified using a normalised difference, (A−P)/(A+P+1)(A−P)/(A+P+1), which is robust to zero counts and downweighs asymmetry estimates at very low expression levels. Asterisks in figures indicate corresponding statistical significance as it follows: * p < 0.05; ** p < 0.01; *** p < 0.005; **** p < 0.001.

## Acknowledgements

This work was supported by the Wellcome Trust [219448/Z/19/Z]. We thank Dimitris Katsanos, Ken Liu, Manish Grover and Simon Berger for experimental, data analysis and bioinformatic support. We want to thank David Matus for providing the HDA- 1::GFP knock-in strain. Some strains were provided by the CGC, which is funded by NIH Office of Research Infrastructure Programs (P40 OD010440). We also thank the facility for Imaging by Light Microscopy (FILM) at Imperial College London, which is part-supported by funding from the Wellcome Trust [104931/Z/14/Z] and the BBSRC [BB/T017929/1].

## Competing Interest Statement

The authors declare no competing interests.

## Author Contributions

MFM performed the majority of experiments and data analysis. BGV, MH, LN, AL, JH, SE contributed to experimental work. MB supervised the study. MFM and MB wrote the manuscript.

## Funding

This work was supported by the Wellcome Trust [219448/Z/19/Z] and a studentship by ‘La Caixa’ Foundation. Open Access funding provided by Imperial College London.

## Data availability

The raw TaDa sequence files and processed signal and peak files have been deposited in the National Center for Biotechnology Information Gene Expression Omnibus under accession number GSE319833.

## Contact for Reagent and Resource Sharing

Further information and requests for resources and reagents should be directed to and will be fulfilled by the Lead Contact, Michalis Barkoulas.

---

**Figure S1:**
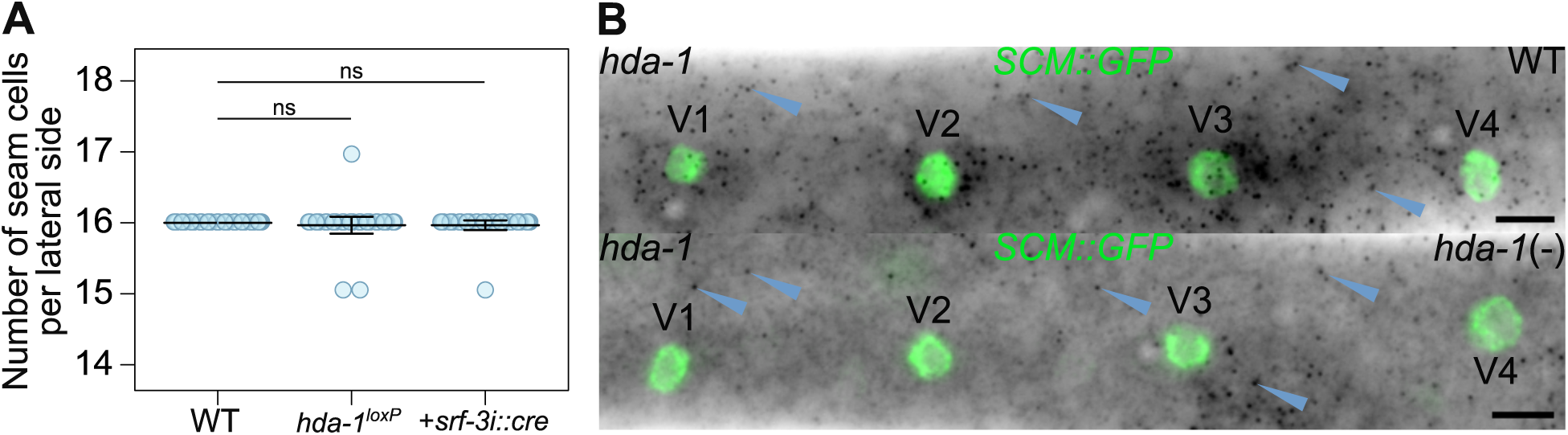
Generation of tissue-specific *hda-1* mutants. **(A)** Seam cell counts in wild-type, animals with *hda-1* locus floxed and animals expressing the CRE recombinase in the seam cells (*+srf-3::cre*). Error bars show the mean ± standard deviation and n = 30 animals per condition. **(B)** Representative images of *hda-1* smFISH at late L1 stage in wild-type (WT) and in *hda-1* mutant animals (*hda-1*(-)). *hda-1*(-) animals don’t show *hda-1* mRNA spots in the seam cells. Blue arrowheads point to *hda-1* mRNA spots in the hypodermis. Seam cell nuclei are labelled using *SCM::GFP* and scale bars are 5 μm.

**Figure S2:**
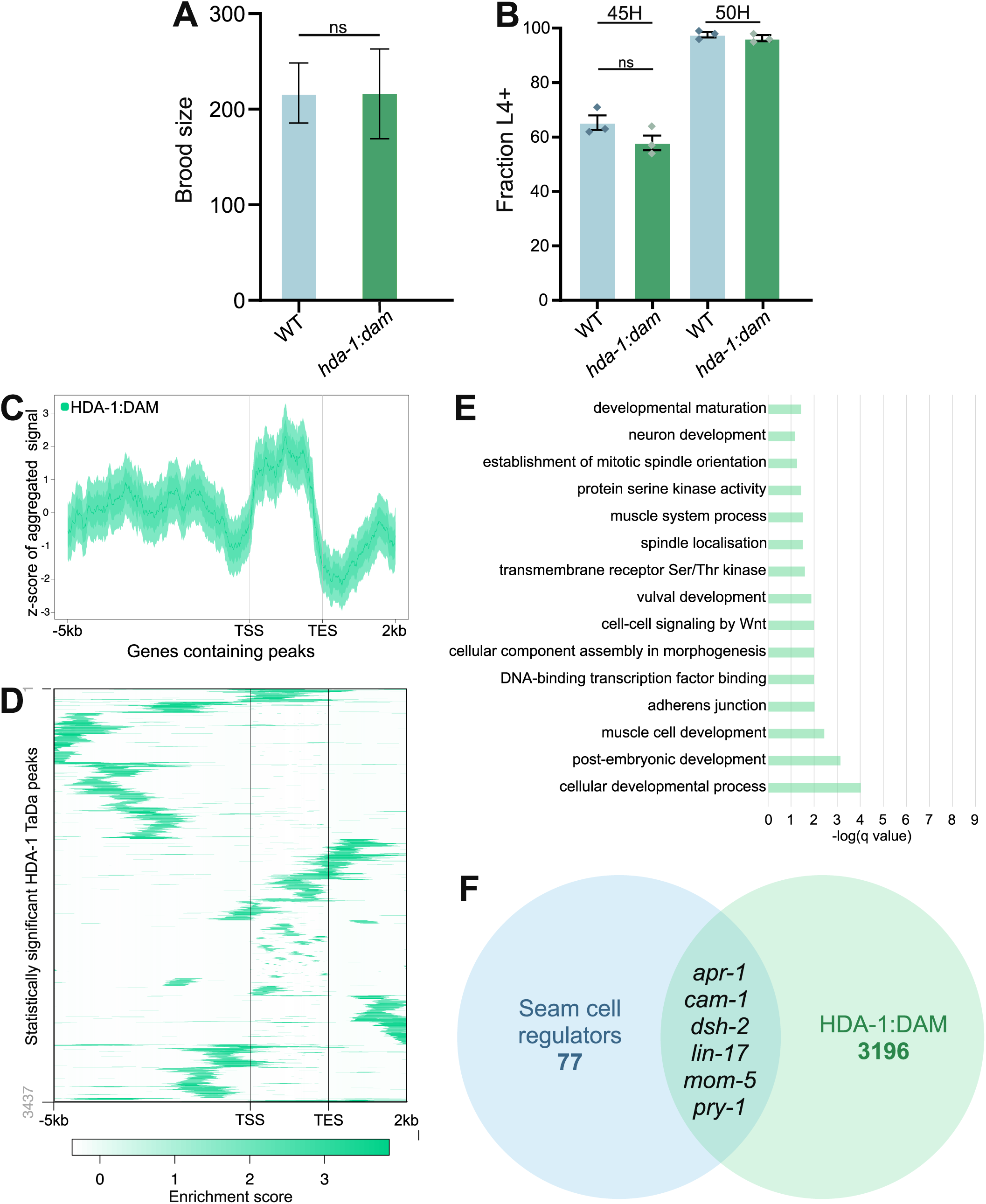
TaDa analysis of HDA-1 binding. **(A)** Quantification of brood size for animals containing single-copy *hda-1:dam* transgene shows no differences compared to control animals, n = 10 animals. **(B)** Quantification of developmental pace by measuring the fraction of animals at or beyond the L4 stage (+L4 stage) at 45h and 50h post egg layering for animals containing a single-copy *hda-1:dam* transgene. Error bars indicate the standard error of proportion, and the bar plot corresponds to the average proportion across three biological replicates. The three datapoints represent values for the three individual replicates. No statistically significant difference in the average proportion of +L4 stage was observed in *hda-1:dam* animals compared to WT with a Chi-square test. **(C)** Aggregation plots of profiling data generated by HDA-1 TaDa showing average enrichment scores in 10-bp bins for regions of equal length across all genes containing statistically significant peaks. Plots show 5 kb upstream of the transcription start site (TSS) and 2 kb downstream of the transcription end site (TES). Gene bodies are condensed into a 2 kb pseudo-length. Shaded areas represent 95% confidence intervals. **(D)** Heatmaps representing the hierarchically clustered localisation and enrichment score of all statistically significant peaks within 5 kb upstream and 2 kb downstream of genes containing peaks. **(E)** Plots of selected significantly enriched terms from GO-term analysis for HDA-1 TaDa. **(F)** Venn diagram highlighting the 6 genes we focused on in this study which are part of the overlap (29 genes) between HDA-1 putative target genes identified with TaDa (green) and a list of 77 genes known to participate in the regulation of seam cell development based on the literature (blue). For a complete list of genes see Table S2.

**Figure S3.**
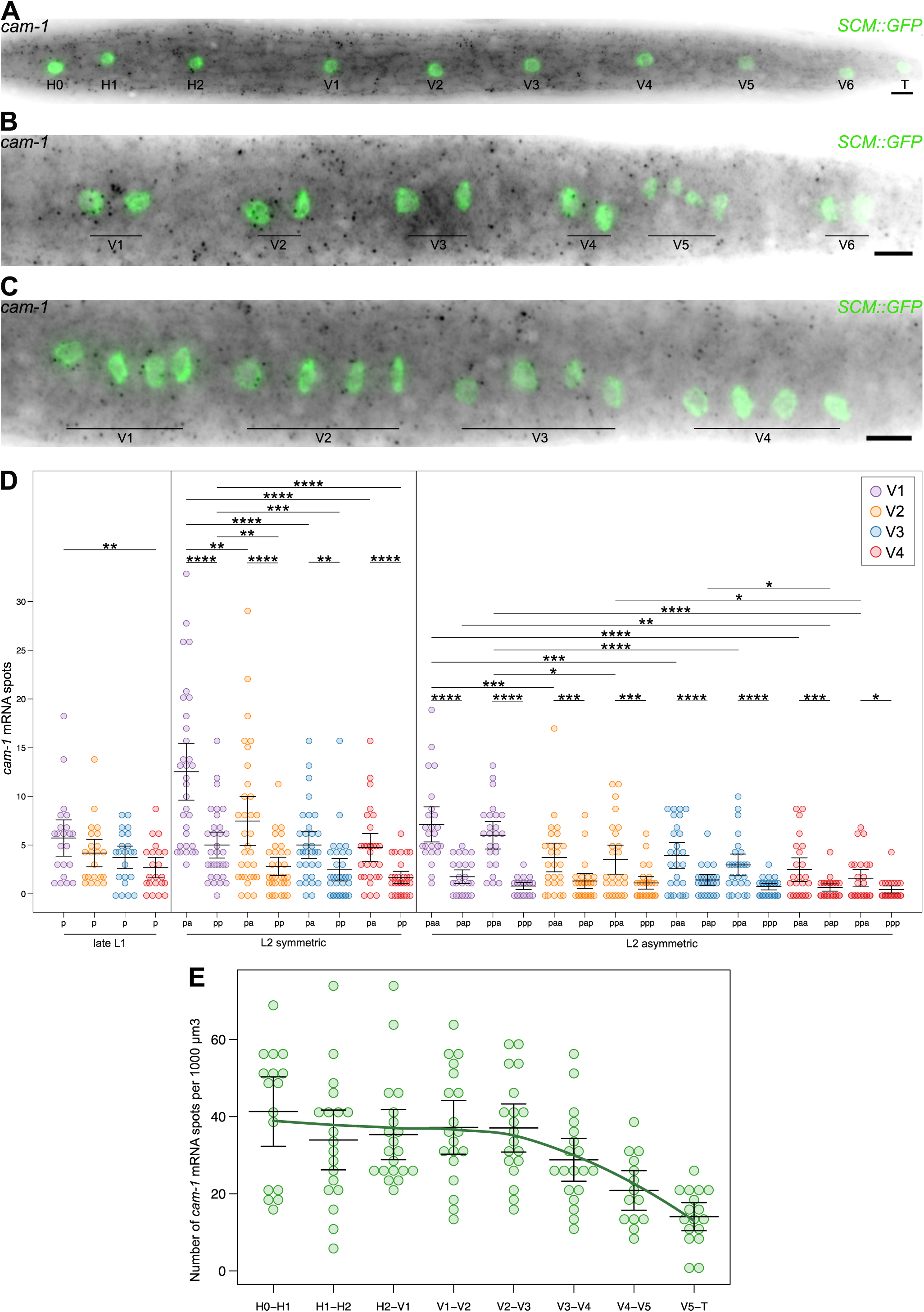
Detailed analysis of *cam-1* expression in wild-type epidermis. (A-C) Representative *cam-1* smFISH images at late L1 (A), and following the L2 symmetric and L2 asymmetric (C) divisions. Seam cell nuclei are labelled with *SCM::GFP* and scale bars are 5μm. **(D)** Quantification of *cam-1* mRNA spots in V1-V4 seam cells before (p) and after the L2 symmetric division (pa, pp), and following the L2 asymmetric division (paa, pap, ppa, ppp); 17 ≤ n ≤ 34 cells per condition. **(E)** Quantification of *cam-1* smFISH spot density in the lateral epidermis across the anterior-posterior axis; n = 21 animals. The solid line represents a LOESS-smoothed trend of mean mRNA expression across the anterior-posterior axis. Error bars in D and E show the mean ± standard deviation. For clarity, in D only statistically significant comparisons are indicated and **** represent *p*<0.001, *** *p*<0.005, ** *p*<0.01 and * *p*<0.05 with a two-tailed t-test.

**Figure S4.**
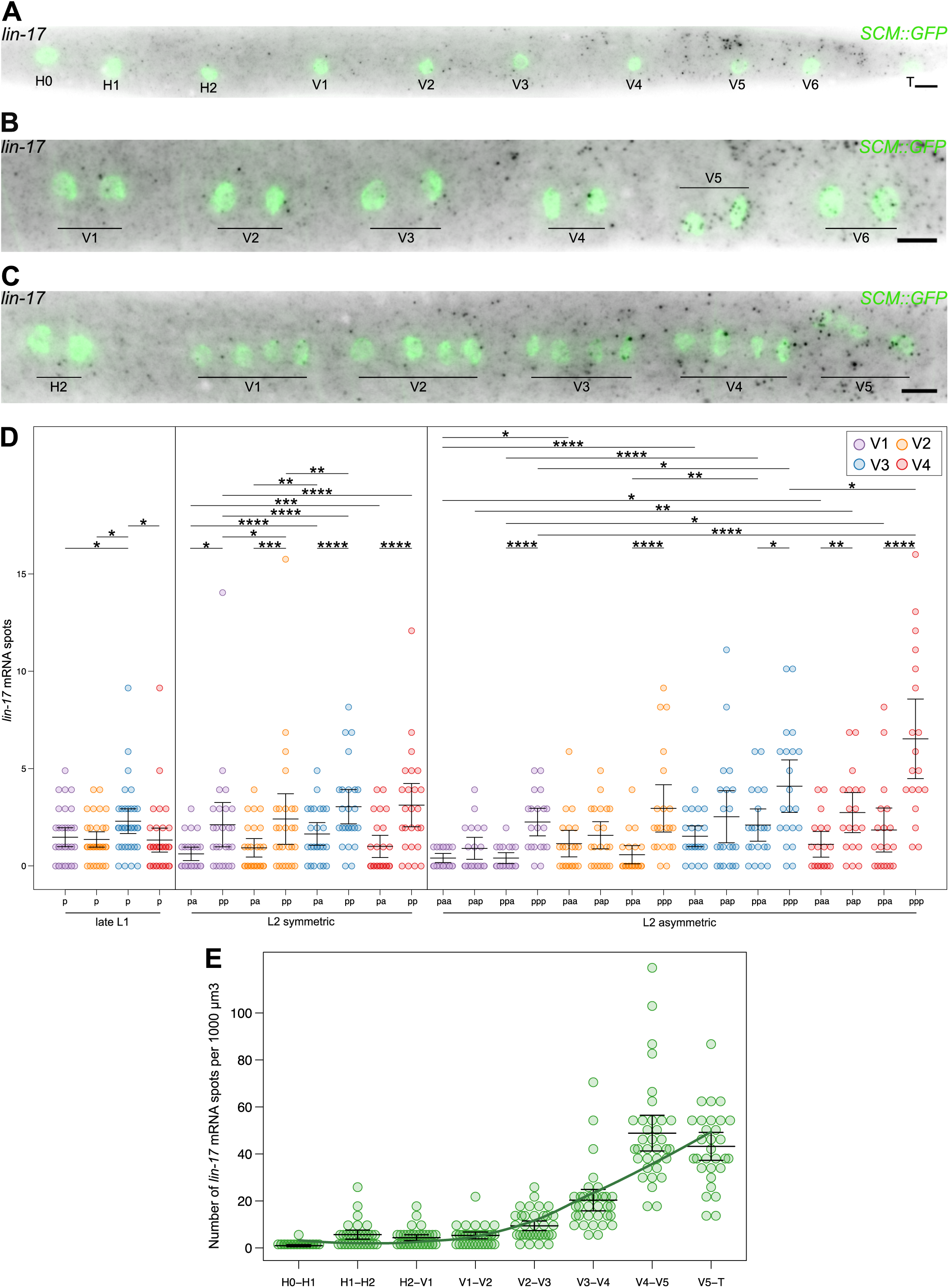
Detailed analysis of *lin-17* expression in wild-type epidermis. (A-C) Representative *lin-17* smFISH images at late L1 (A), and following the L2 symmetric (B) and L2 asymmetric (C) divisions. Seam cell nuclei are labelled with *SCM::GFP* and scale bars are 5μm. **(D)** Quantification of *lin-17* mRNA spots in V1-V4 seam cells before (p) and after the L2 symmetric division (pa, pp), and following the L2 asymmetric division (paa, pap, ppa, ppp); 20 ≤ n ≤ 34 cells per condition. **(E)** Quantification of *lin-17* smFISH spot density in the lateral epidermis across the anterior-posterior axis; n = 33 animals. The solid line represents a LOESS-smoothed trend of mean mRNA expression across the anterior-posterior axis. Error bars in D and E show the mean ± standard deviation. For clarity, in D only statistically significant comparisons are indicated and **** represent *p*<0.001, *** *p*<0.005, ** *p*<0.01 and * *p*<0.05 with a two-tailed t-test.

**Figure S5:**
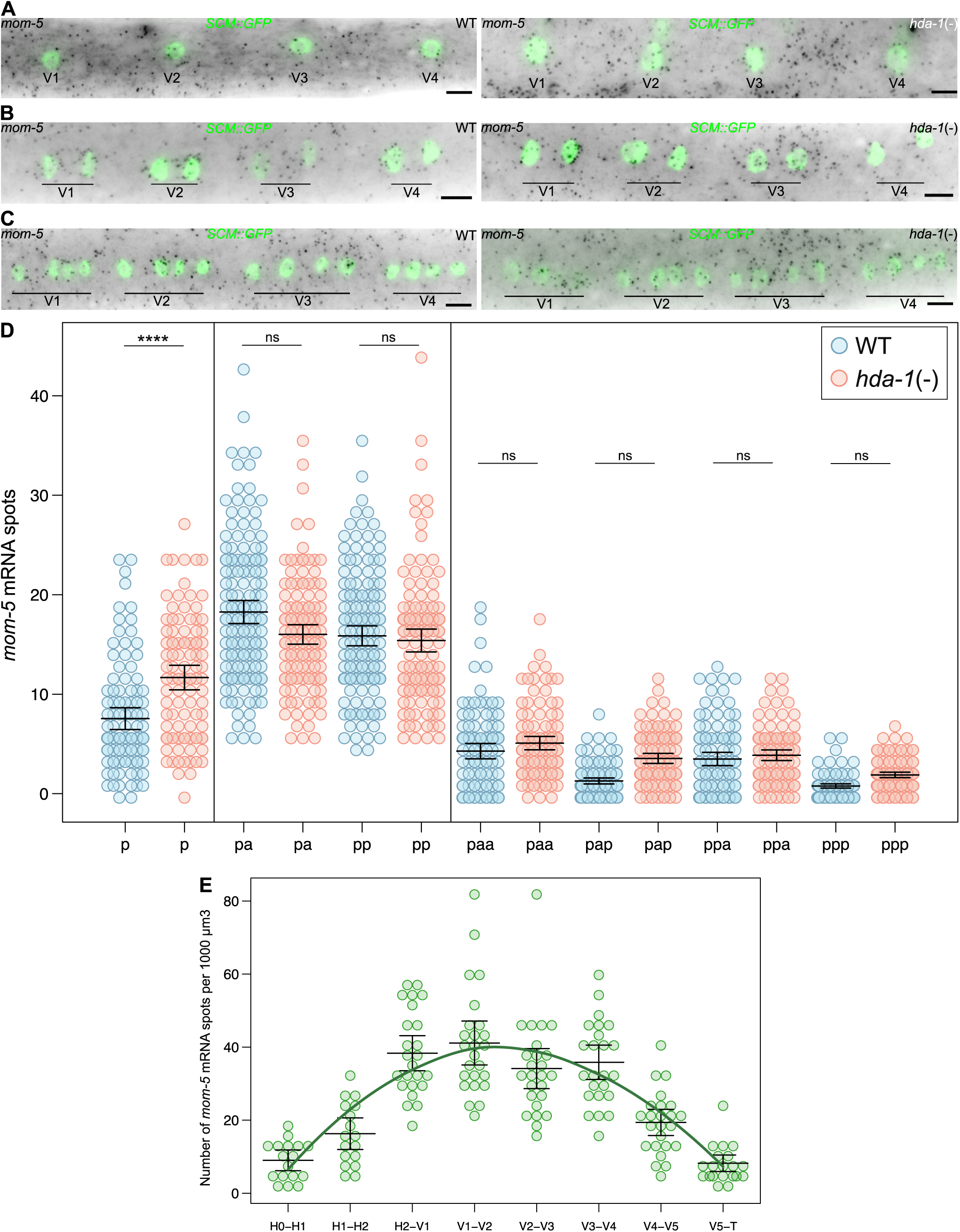
Analysis of *mom-5* expression in *hda-1* mutants. (A-C) Representative *mom-5* smFISH images of WT and tissue-specific *hda-1* mutant animals (*hda-1*(-)) at late L1 (A) and following the L2 symmetric (B) and L2 asymmetric divisions (C). Seam cell nuclei are labelled using *SCM::GFP* and scale bars are 5 μm. **(D)** Quantification of *mom-5* mRNA spots in V1-V4 lineages in WT (blue) and *hda-1* mutant animals (red) at late L1 (p) and following the L2 symmetric (pa, pp) and L2 asymmetric divisions (paa, pap, ppa, ppp); 98 ≤ n ≤ 99 (p), 149 ≤ n ≤ 125 (pa,pp) and 87 ≤ n ≤ 102 cells per condition cells. **(E)** Quantification of *mom-5* smFISH spot density in the lateral epidermis across the anterior-posterior axis; n = 23 animals. The solid line represents a LOESS-smoothed trend of mean mRNA expression across the anterior-posterior axis. Error bars show the mean ± standard deviation and **** represent *p*<0.001 with a two-tailed t-test.

**Figure S6:**
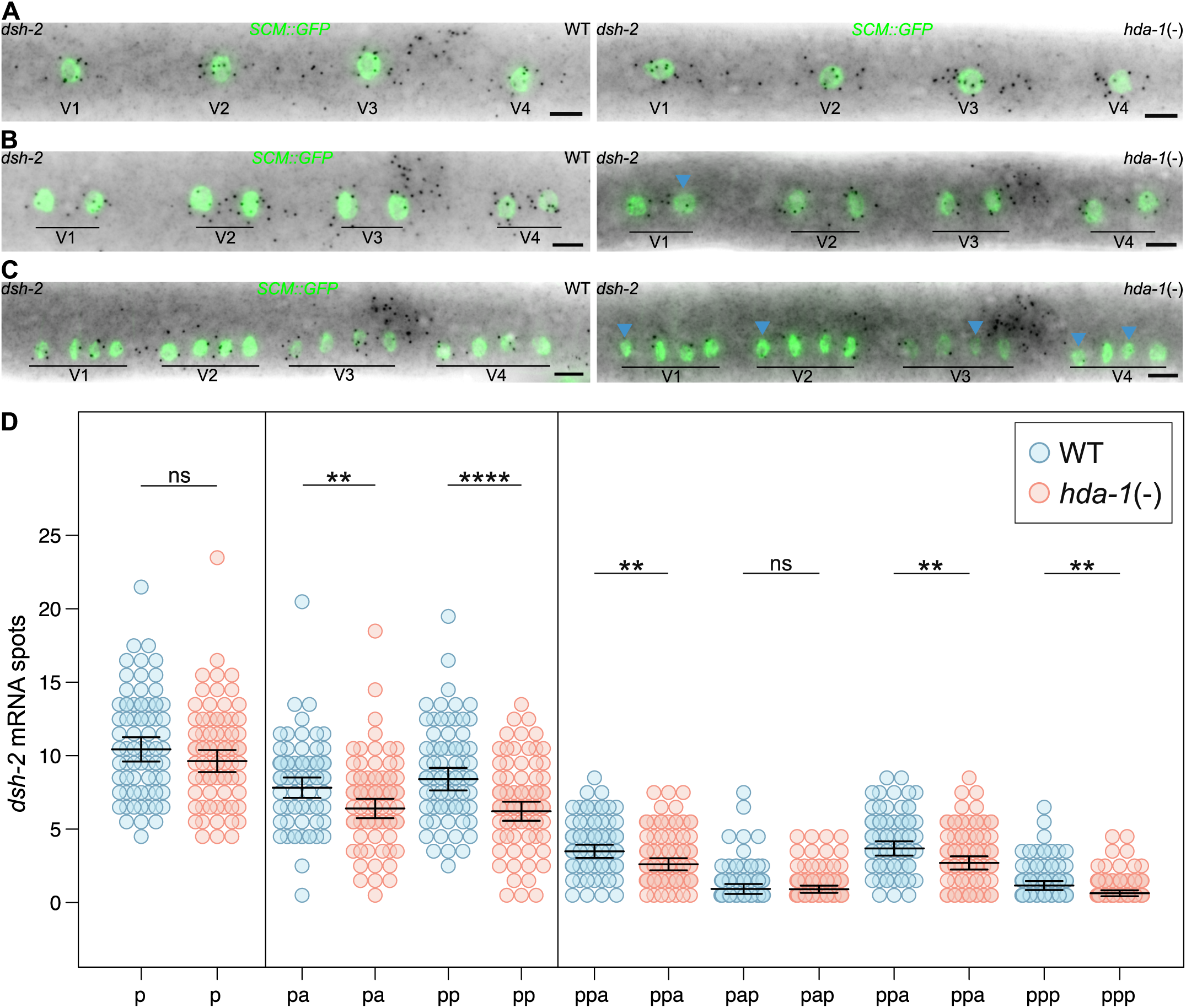
Analysis of *dsh-2* expression in *hda-1* mutants. (A-C) Representative *dsh-2* smFISH images of WT and tissue-specific *hda-1* mutant animals (*hda-1*(-)) at late L1 (A) and following the L2 symmetric (B) and L2 asymmetric divisions (C). Seam cell nuclei are labelled using *SCM::GFP* and scale bars are 5 μm. **(D)** Quantification of *dsh-2* mRNA spots in V1-V4 lineages in WT (blue) and *hda-1* mutant animals (red) at late L1 (p) and following the L2 symmetric (pa, pp) and L2 asymmetric divisions (paa, pap, ppa, ppp); 72 ≤ n ≤ 76 (p), 72 ≤ n ≤ 79 (pa,pp) and 70 ≤ n ≤ 79 cells per condition cells. Error bars show the mean ± standard deviation and ** represent *p*<0.01 and **** represent *p*<0.001 with a two-tailed t-test.

**Figure S7:**
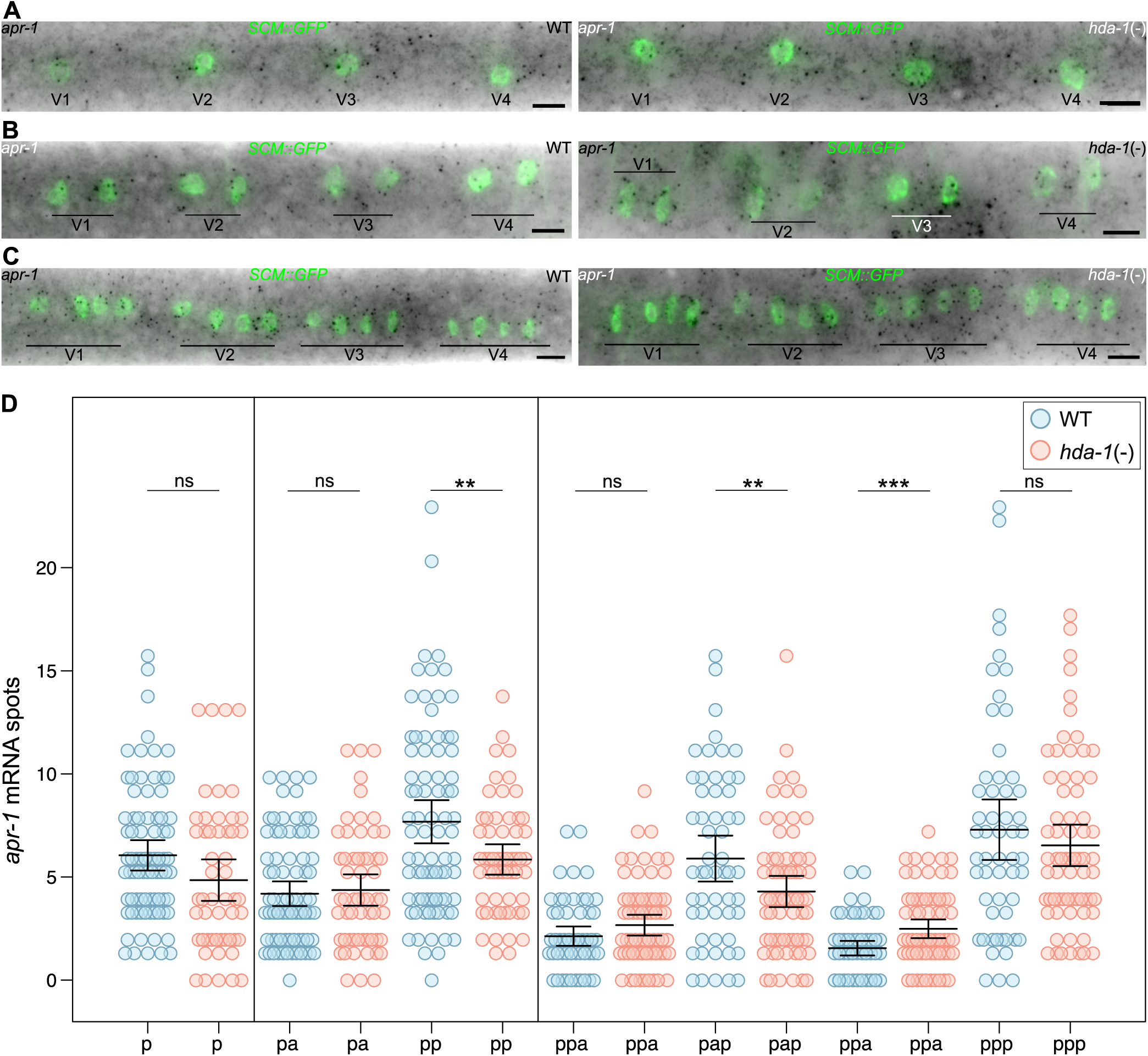
Analysis of *apr-1* expression in *hda-1* mutants. (A-C) Representative *apr-1* smFISH images of WT and tissue-specific *hda-1* mutant animals (*hda-1*(-)) at late L1 (A) and following the L2 symmetric (B) and L2 asymmetric divisions (C). Seam cell nuclei are labelled using *SCM::GFP* and scale bars are 5 μm. **(D)** Quantification of *apr-1* mRNA spots in V1-V4 lineages in WT (blue) and *hda-1* mutant animals (red) at late L1 (p) and following the L2 symmetric (pa, pp) and L2 asymmetric divisions (paa, pap, ppa, ppp); 52 ≤ n ≤ 80 (p), 58 ≤ n ≤ 80 (pa,pp) and 55 ≤ n ≤ 68 cells per condition cells. Error bars show the mean ± standard deviation and ** represent *p*<0.01 and *** represent *p*<0.005 with a two-tailed t-test.

**Figure S8:**
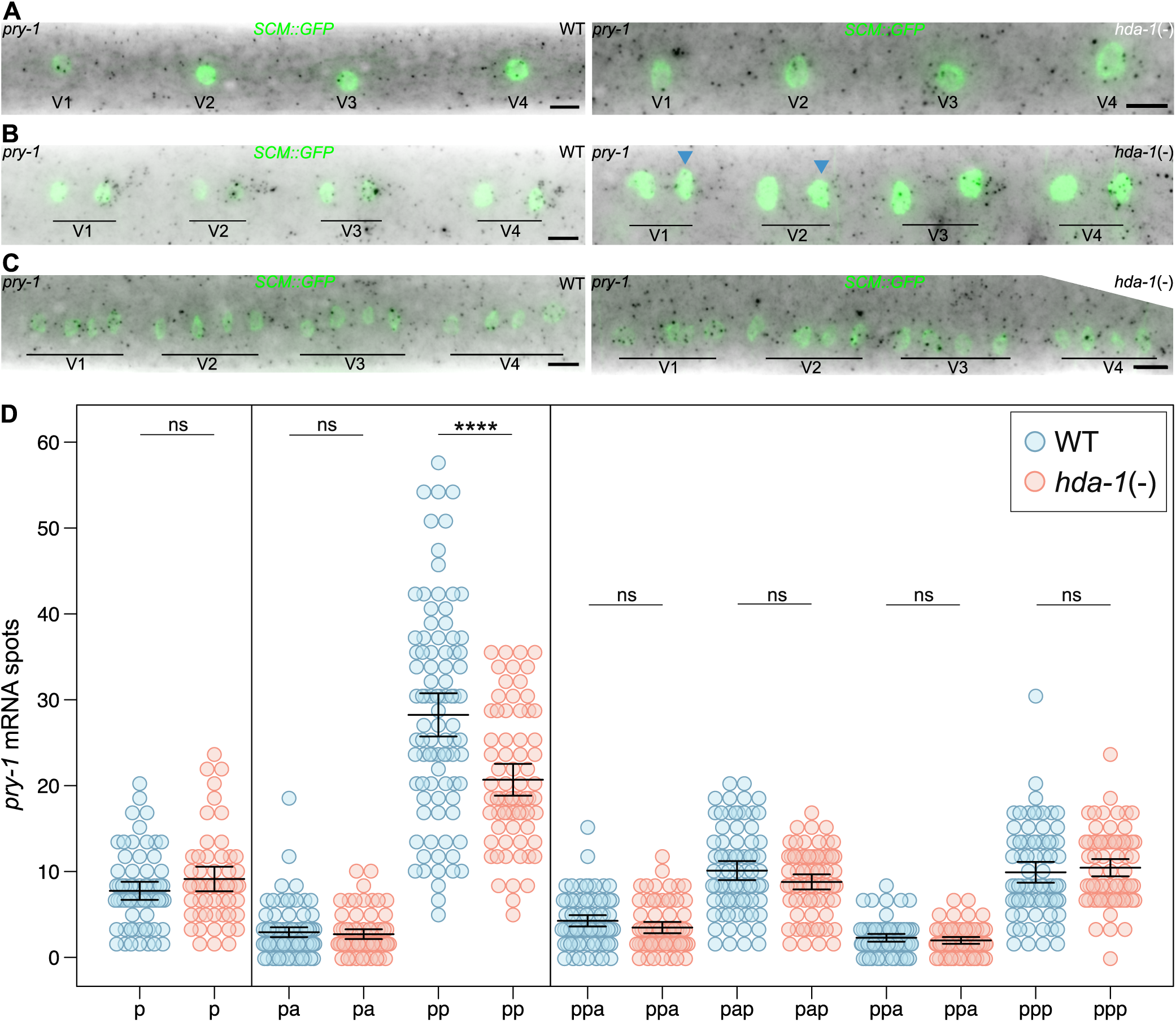
Analysis of *pry-1* expression in *hda-1* mutants. (A-C) Representative *pry-1* smFISH images of WT and tissue-specific *hda-1* mutant animals (*hda-1*(-)) at late L1 (A) and following the L2 symmetric (B) and L2 asymmetric divisions (C). Seam cell nuclei are labelled using *SCM::GFP* and scale bars are 5 μm. **(D)** Quantification of *pry-1* mRNA spots in V1-V4 lineages in WT (blue) and *hda-1* mutant animals (red) at late L1 (p) and following the L2 symmetric (pa, pp) and L2 asymmetric divisions (paa, pap, ppa, ppp); 54 ≤ n ≤ 67 (p), 70 ≤ n ≤ 91 (pa,pp) and 70 ≤ n ≤ 75 cells per condition cells. Error bars show the mean ± standard deviation and **** represent *p*<0.001 with a two-tailed t-test.

## Notes

https://www.ncbi.nlm.nih.gov/geo/query/acc.cgi?acc=GSE319833

### Competing Interest Statement

The authors have declared no competing interest.

